# Tubulin glycylation regulates microtubule-protein interactions that are key for ciliary stability and trafficking

**DOI:** 10.64898/2026.03.26.714406

**Authors:** Sanjana Mullick, Chaya Suresh Kumar, Sucheta Dey, Prajwal B Koushik, Rayees Ganie, Susobhan Mahanty, Minhajuddin Sirajuddin, Sudarshan Gadadhar

**Affiliations:** Institute for Stem Cell Science and Regenerative Medicine (iBRIC-inStem), GKVK Campus, Bellary Road, Bangalore - 560065, Karnataka, India; Regional Centre for Biotechnology (RCB), NCR Biotech Science Cluster, 3rd Milestone, Faridabad-Gurugram Expressway, Faridabad, 121001, Haryana (NCR Delhi), India; Manipal Academy of Higher Education University, Karnataka, 576104, India

**Keywords:** tubulin glycylation, tubulin posttranslational modifications, molecular motors, microtubule-associated proteins, kinesin, cilia

## Abstract

Tubulin glycylation, a cilia-specific posttranslational modification is emerging as a potentially key regulator of ciliary axonemal microtubules. However, insights into the functional consequences of glycylation have remained limited. Here, using *in vitro* reconstitution assays with unmodified or custom-glycylated tubulin, we provide a systematic mechanistic analysis of glycylation-dependent regulation of motors and microtubule-associated proteins. Our studies highlight that glycylation selectively enhances ciliary kinesin-2 motility while reducing kinesin-1 activity, suggesting a role in promoting efficient intraflagellar transport along axonemal microtubules. Moreover, glycylation protects microtubules from decay by suppressing the activities of the depolymerase MCAK and severing enzyme spastin, thereby enhancing stability. Notably, this regulation is dependent on the proportion of glycylation on the microtubule surface, coupled with concomitant reduction of glutamylation. Thus, by generating microtubule surfaces with distinct biochemical states, we establish that combinatorial modification patterns define functional microtubule properties especially in cilia. Together, our findings provide the first comprehensive mechanistic framework for tubulin glycylation in regulating molecular motors and MAPs in cilia, establishing glycylation as a key determinant of motor selectivity and microtubule stability within the axoneme.

## Introduction

Microtubules perform a plethora of functions within the eukaryotic cytoskeleton, and a major contributor to this functional heterogeneity is the tubulin code ^1,2^. Mammals have 9 α- and 9 β-tubulin isotypes (https://www.genenames.org/data/genegroup/#!/group/778) differentially expressed in a cell and tissue-specific manner, where the isotype diversity majorly resides on the unstructured Carboxy-terminal tails (CTT). Both α- and β-tubulin undergo diverse tubulin posttranslational modifications (PTMs; reviewed in ^1,2^). Apart from phosphorylation, methylation and acetylation that occur on the core globular structure of tubulin, all other PTMs are confined to the CTTs ^1–3^.

The tubulin code fine-tunes microtubule-associated protein (MAP) interactions, specifically those governed by tubulin PTMs, based on the location and pattern of modifications. Axoneme, the microtubule-based ciliary core is one of the diverse specialised microtubules enriched with many of these PTMs. Cilia and flagella regulate myriad cellular functions and hence organ physiology and the axonemes are conserved structurally across evolution. The microtubules forming the axonemes are enriched in two key tubulin modifications, glutamylation and glycylation, with glycylation uniquely observed so far only on cilia and flagella ^3–7^. Both these modifications are catalysed by tubulin tyrosine-ligase like (TTLL) family of enzymes that have both substrate (α- or β-tubulin) and reaction specificities (mono- vs poly-modification; reviewed in ^2,3^) and compete for the same modification sites on both α- and β-tubulin. So far, glutamylation has been demonstrated to fine-tune microtubule-MAP interactions using *in vitro* analyses, where different kinesin motors exhibited differential sensitivity to microtubules with regulated patterns of glutamylation^8^. Spastin and katanin are also similarly regulated either by the length of the glutamate chain ^9,10^ or their specificity for either α- or β-tubulin tails ^11^.

Glycylation, on the other hand is the most understudied and underappreciated modification. It is involved in controlling motility in different model organisms ^12–16^ as well as stabilizing the axonemes and regulating primary cilia length^17^. Its importance in physiology is highlighted by the fact that absence of this modification can result in ciliopathies like male subfertility^13^, retinal degeneration ^18^ and colorectal cancer ^19^. However, an in-depth molecular understanding of the regulation of ciliary microtubule-MAP interactions by glycylation is missing. So far, the only available evidence of its regulation of motors and MAPs is of its control of axonemal dyneins in sperm ^13^, established through structural analyses and its antagonism to glutamylation in regulating katanin activity *in vitro* ^11^. Cilia and flagella biogenesis, maintenance, and functions depend on intraflagellar transport (IFT), which helps in carrying the cargoes into and out of the cilia using molecular motors, kinesins and dyneins ^20–22^, as well as the MAPs that maintain ciliary length all of which intricately interact with axonemal microtubules. Glycylation, being exclusive to the axonemes of cilia and flagella, can potentially regulate these interactions, and understanding how this is achieved can provide key molecular insights to the importance of glycylation in ciliated cells.

Isolating axonemal microtubules to understand glycylation and its regulation of microtubule-protein interactions is challenging, and the amount of tubulin obtained is limiting even for *in vitro* analyses. Hence, here we devised an approach to generate homogeneous, unmodified or custom-glycylated tubulin and tested it against the most important readers of the tubulin code, the molecular motors and microtubule-stabilising MAPs ^1,23,24^. Our studies show that glycylation differentially affects kinesin-1 and kinesin-2, with gliding velocities of glycylated MT on kinesin-1 slower than on kinesin-2, independent of their binding affinities to these MTs. The specific enhancement of kinesin-2 motor activity emphasises the impact of glycylation in cilia as kinesin-2 is a motor involved in ciliary intraflagellar transport (IFT) and hence, glycylation may provide a favourable molecular surface for IFT within the cilia. Furthermore, we also show that glycylation reduces the kinetics of spastin and the depolymerising mitotic centromere-associated kinesin (MCAK; kinesin-13). While loss of glycylation was shown to impact axonemal dynein interactions, and thus sperm motility defects^13^, our *in vitro* study establishes that increased glycylation also perturbs motor and MAP interactions. Hence, this establishes that an optimal level of both glycylation and glutamylation is essential for fine regulation of ciliary MAPs and optimal motor activity. Any deviations from these levels can perturb these interactions, thus impacting ciliary functions.

## Results

### Development of a toolbox of custom-modified tubulins and molecular motors

Most *in vitro* microtubule-MAP studies utilise the purified brain tubulin, which is highly enriched in diverse tubulin PTMs, especially glutamylation. While this acts as our highly glutamylated tubulin fraction in our assays, it was also an internal control to assess the veracity of our assays. Since we wanted to establish the impact of glycylation specifically, we needed microtubules that lacked either glutamylation or glycylation (unmodified), as well as microtubules with only glycylation. Our need for such unmodified and tubulin custom-modified with only glycylation was fulfilled by purifying them from Lenti-X^TM^ 293T (Lenti-X) cells (Fig. S1A). Microtubules in Lenti-X cells undergo very low levels of glutamylation and glycylation ^17,25^ and hence, serve as an excellent source for unmodified tubulin. A stable Lenti-X cell line expressing mCherry-TTLL3 was engineered to generate the custom-modified glycylated tubulin (*see methods for details)*. Both the Lenti-X and Lenti-X-mCherry-TTLL3 cells were grown as suspension cultures for purifying adequate amounts of tubulin. Purification of all the different tubulins was carried out using repeated cycles of polymerisation at 30 °C and depolymerisation on ice at 4 °C^25^ (*see methods for details*). The purity was assessed using SDS-PAGE and Coomassie blue staining (Fig. S1B-D). Every purification cycle yields ∼300 mg brain tubulin, ∼600 μg of unmodified tubulin and ∼500 µg of glycylated tubulin. Immunoblot analyses of each of these purified tubulins revealed a distinct pattern, which matched our expectations (Fig. S1E). The glycylated tubulin we purified also showed increased glycylation predominantly on β-tubulin with some degree of glycylation on α-tubulin, since TTLL3 preferentially modifies β-tubulin over α-tubulin ^26^.

In the study we tested two different cargo-carrying motors, human kinesin-1 (KIF5B) and *C. elegans* kinesin-2 (Osm3) based on their well-characterised motility properties *in vitro* ^27,28^. The truncated dimeric versions of these motors were used in our assays (*see methods for details)*. We also tested the depolymerising kinesin-13, MCAK and severing protein, spastin both implicated to have roles in primary cilia ^29,30^. All these were expressed and purified from bacteria either as 10x-His tag proteins or as GST tag proteins. The purity of the proteins was determined by SDS-PAGE and Coomassie blue staining (Fig. S1F). Kinesin-1 is a multimeric motor and the presence of multiple bands in the purified protein (Fig. S1F) likely reflects different forms of the motor expressed in the bacterial system. To enhance the active motor fraction, the purified motors were additionally enriched through a microtubule bind-and-release step (Fig. S1G-H).

### Increased β-tubulin glycylation reduces the kinetics of kinesin-1, but not kinesin-2

We first tested how the molecular motors behave in presence of MTs either unmodified or enriched with glutamylation or glycylation. Our purified GFP-tagged KIF5B consisted of the N-terminal 560 amino acids, corresponding to the motor domain (K560^31^). We first tested how K560-GFP interacts with brain MTs, unmodified, and glycylated MTs by TIRF-based assessment of MT gliding on anchored K560-GFP motors (Fig. 1A). We observed that among the three microtubules, unmodified MTs showed the fastest gliding with a median velocity of ∼270 nm s^−1^ followed by brain MTs at 184 nm s^−1^, and the glycylated MTs being the slowest at 103 nm s^−1^ (Fig. 1B, C; Fig. S2A, Movie S1, Table 1, Table S1).

**Figure 1:**
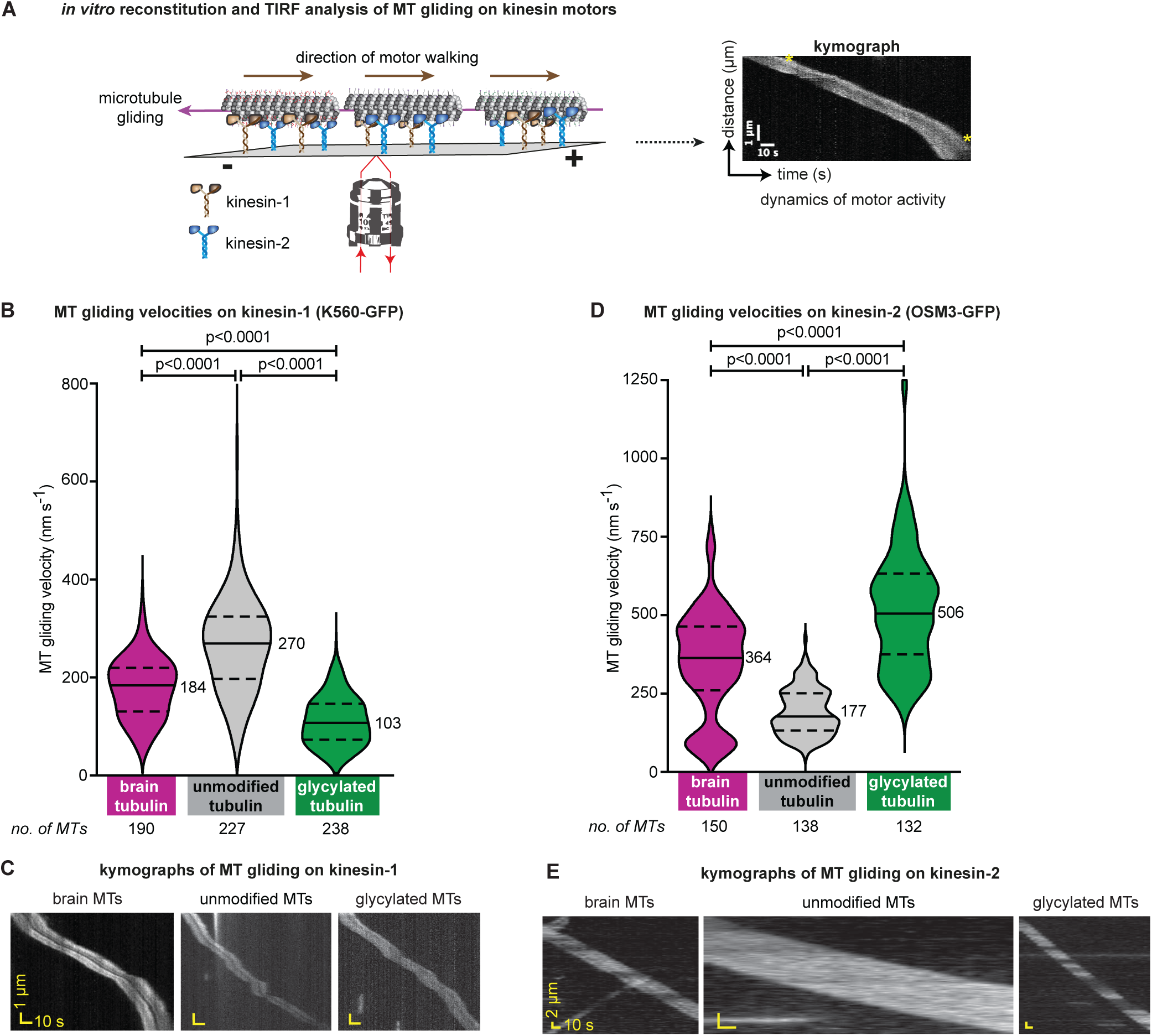
Regulation of kinesin motors by tubulin glycylation. **A.** Schematic depiction of the TIRF microscope-based *in vitro* microtubule gliding of the three MT variants, with either kinesin-1 (K560) or kinesin-2 (Osm3). The motility chambers coated with the respective motors are incubated with the different microtubule variants and the MT gliding is assessed in presence of ATP. The gliding visualized using TIRF microscopy is then analysed by building the kymographs and analysing motor velocity to determine the speed of motor activity on each of the different MTs. The yellow stars indicate the start and the end points of the kymographs, the slope of which was used to calculate the velocity. **B.** *In-vitro* MT gliding assay on K560-GFP shows that the velocity of the K560 is reduced when the MTs are heavily glycylated. The violin plot represents a mean of 3 independent experiments where the numbers and solid line denote the median velocity and dashed lines the 25th and 75th quartiles. The graph shows that the gliding velocities for glycylated MTs reduces by ∼2.6-fold and ∼1.7-fold compared to unmodified and brain MTs. Individual experiments are shown in Fig. S2A. **C.** Representative kymographs of for brain MTs, HEK MTs and glycylated MTs gliding on K560 that were used to quantify the gliding velocity of each MTs. The x-axis depicts time (s) and the y-axis distance of gliding (µm). **D.** *In-vitro* MT gliding assay on Osm3ΔH2-GFP shows that contrary to kinesin-1, the velocity of MT gliding on Osm3 is enhanced when the MTs are glycylated. The violin plot of the gliding velocities shows that the glycylated MTs had a median velocity of 506 nm s^−1^, which is ∼1.4-fold higher than brain MTs (∼364 nm s^−1^) and ∼2.85-fold higher than unmodified MTs (∼177 nm s^−1^). The graph represents a mean (± SEM) of 3 independent experiments with the numbers and solid line denoting the median value and the dashed lines, the 25th and 75th quartiles. Individual experimental replicates are shown in Fig. S2B. **E.** Representative kymographs of for brain MTs, HEK MTs and glycylated MTs gliding on Osm3ΔH2 that were used to quantify the gliding velocity of each MTs. The slope of the kymographs indicates the velocities with the x-axis depicting time (s) and y-axis, the distance of gliding (µm).

**Table 1:** Kinetic parameters of the different motors and MAPs on MTs with diverse PTM patterns. Values representing the gliding velocities, depolymerisation rates and severing activities are the mean (± SEM) and n represent the total number of measurements. For kinesin-1 and kinesin-2, the n represents number of gliding MTs analysed. For MCAK activity, n is the number of microtubules ends analysed. For the extent of severing by spastin, the n indicates the number of MTs undergoing both severing and depolymerisation analysed, while for the rate of severing, n is the number of MTs showing clear severing events during imaging. For all the proteins, the values are combined from 3 individual biological experiments, wherein for each experiment, the MTs from multiple imaging chambers were analysed.

While kinesin-1 is predominantly a cytoplasmic motor, kinesin-2 is a key player involved in intraflagellar transport within the cilia ^32^, specifically in anterograde IFT ^33^. Glycylation being a modification exclusive to the cilia, we tested how a ciliary motor would interact with microtubules having only glycylation. Our MT gliding assays showed that contrary to what we observed for kinesin-1, the addition of glycine enhanced the median velocity of Osm3ΔH2-GFP at 506 nm s^−1^, with brain MTs gliding at 364 nm s^−1^ similar to what has already been reported earlier ^34^, while the unmodified MTs glide ∼2.85-fold slower at 177 nm s^−1^ (Fig. 1D, E; Fig. S2C Movie S2, Table 1, Table S2). This suggests that kinesin-2 depends on both glutamylation and glycylation for optimum activity, with a slightly higher preference for glycylation.

To further understand whether this difference in activities was driven solely by the motor activity or also influenced by the binding of the motor to the different MTs, we tested the binding of the GFP-tagged motors in our *in vitro* reconstitution system. K560–GFP exhibited differential binding affinities, with the strongest binding to brain-derived MTs and a concentration-dependent loss of binding to glycylated MTs (Fig S2B). Notably, any concentration less than 56 nmol of K560 failed to bind glycylated MTs while maintaining substantial binding to both brain and unmodified MTs. In contrast, Osm3 displayed minimal difference in binding to either brain MTs or glycylated MTs, with much lesser binding observed to unmodified MTs (Fig S2D). These indicate that glycylation may enhance kinesin-2 (Osm3) association with MTs, while counteracting the kinesin-1 binding, which may underlie the differences in the MT gliding observed between the two motors on glycylated MTs.

In summary, tubulin glycylation enhances Osm3 motility, reflected in elevated velocity. The higher binding of Osm3 to glycylated MTs further highlights improved microtubule-motor interactions. Both support a mechanistic role for glycylation in promoting efficient kinesin-2 driven transport along ciliary axonemal microtubules, thus suggesting why robust intraflagellar transport within cilia requires glycylation.

### Tubulin glycylation negatively regulates depolymerising and severing MAPs

Microtubules undergo constant polymerisation and depolymerisation, which help regulate various aspects of cellular dynamics. While this dynamics is slower within the cilia, the axonemal MTs also show regulation of their length by the depolymerising and severing MAPs. Since glycylation is suggested to stabilise the cilia ^17,35^ and increased glycylation leads to increased cilia length ^17^, we tested the impact of glycylation on both the depolymerising and severing MAPs. Among these, the predominant ones regulating cilia dynamics are the kinesin-13 family motors and the severing enzymes spastin and katanin ^30,36–39^.

Within the kinesin-13 family motors, MCAK is well characterised ^40,41^ and the predominant depolymeriser implicated with a key role in ciliogenesis, where excessive accumulation of MCAK at the basal bodies can negatively impact cilia formation ^37^. We tested the rate of depolymerisation induced by MCAK on the three microtubule variants (Fig. 2A). Our analyses showed that glycylated MTs were depolymerised much slower than brain MTs (Fig. 2B,C, Movie S3). When the depolymerisation rates were quantified, we observed that the median depolymerisation rate of the glycylated MTs, was at 0.08 μm min^−1^ which is lower than unmodified MTs, which depolymerised at 0.22 μm min^−1^. The highly glutamylated brain MTs depolymerised at 0.33 μm min^−1^, which matches what has been reported earlier ^42^. Thus, the glycylated MTs were depolymerizing at ∼2.75-fold lower than unmodified MTs and ∼4-fold lower rate than brain MTs (Fig. 2D Fig. S3, Table 1, Table S3). The distribution plot further shows that most of the glycylated MTs were depolymerised by MCAK at ≤ 0.1 μm min^−1,^ whereas brain MTs were depolymerised at 0.3-0.4 μm min^−1,^ with many having rates ≥ 0.5 μm min^−1^ (Fig. 2E, Table S3). Though MCAK favours the plus-end of the MTs, it can depolymerise from both ends in an *in vitro* assay ^43^. We observed similar results with the brain MTs and to some degree, even the unmodified MTs, while with the glycylated MTs, the depolymerisation was predominantly confined to one end of the MTs (Fig. 2C, Movie S3), because the overall rate of MCAK activity is much slower on glycylated MTs.

**Figure 2:**
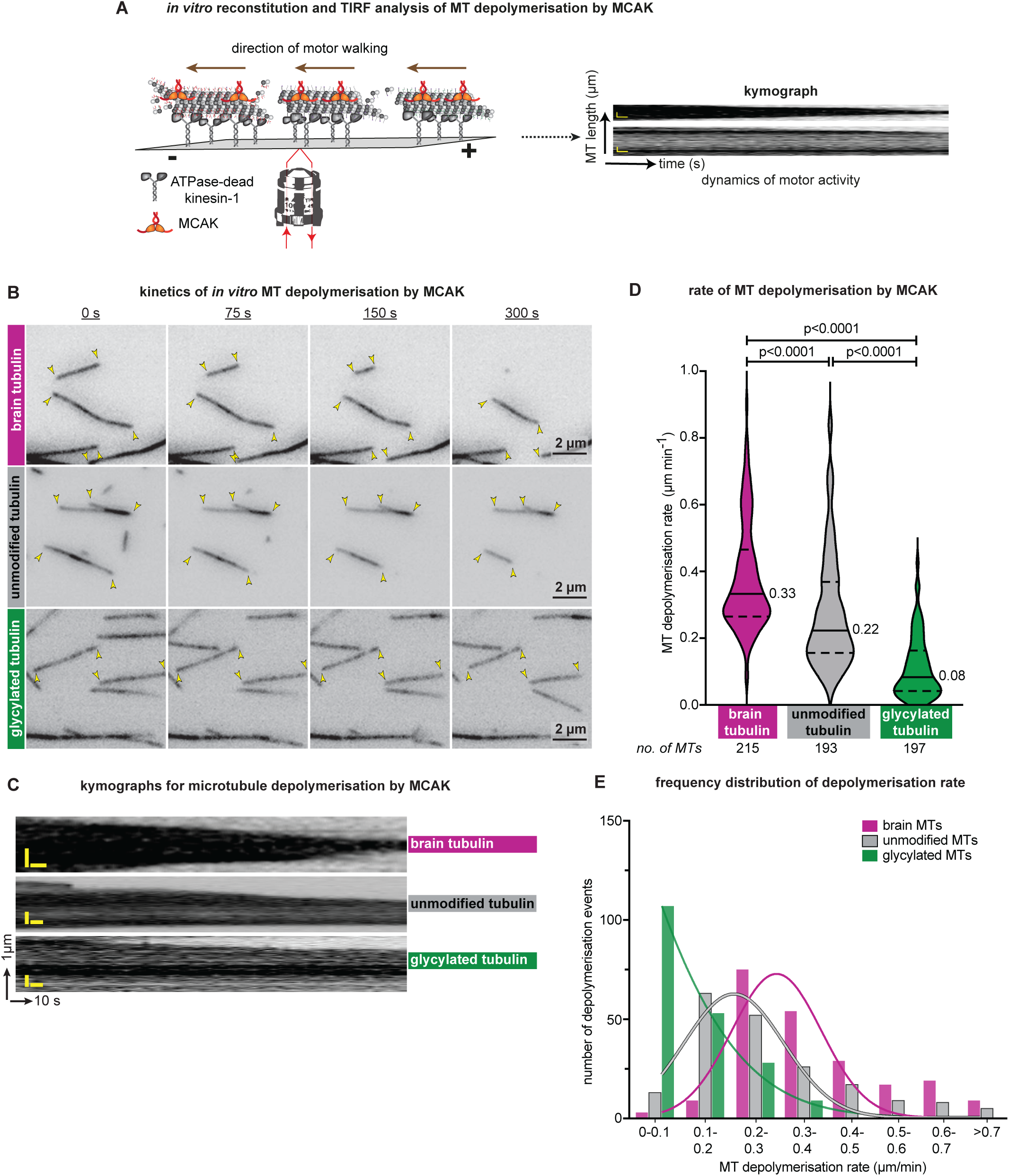
Microtubule depolymerisation by kinesin-13 (MCAK) is reduced on glycylated MTs. **A.** Schematic depiction of the TIRF microscope-based *in vitro* microtubule depolymerisation assay with kinesin-13 (MCAK). The motility chambers coated with ATPase-dead Kif5B are incubated with the different PTM variant MTs and subsequently flushed with the motility mix consisting of ATP and 25nM MCAK. The MT depolymerisation is imaged using the 100x TIRF objective and the videos analysed, kymographs established and the rate of MT depolymerisation determined from these kymographs. **B.** Snapshots of the kinetics of MCAK-induced depolymerisation of the different MTs. From the still images, we observe that brain MTs begin to depolymerise by 75s, and by 300 s the length has reduced by ≥50%. The glycylated MTs, on the other hand remain substantially intact even after 300 s and there is only a marginal change in their length, predominantly from only one end of the MTs. Unmodified MTs show that they are in a state in between the two modified versions of MTs. Yellow arrowheads point to the two ends of the microtubule that are undergoing depolymerisation. **C.** Representative kymographs obtained from the videos of MT depolymerisation show that the brain MTs have a rapid reduction in their length with time, with both the ends depolymerising almost at similar rates. Unmodified tubulin also undergoes a quick depolymerisation. Glycylated MTs show reduced levels of depolymerisation, but only from one of the ends of the MTs. **D.** Violin plot of the rate of MT depolymerisation of the different MT variants quantified from the *in vitro* reconstitution assays. The graph shows that the median rate of MCAK-induced MT depolymerisation for glycylated MTs was 0.08 µm min^−1^ which is ∼4-fold lower than brain MTs (0.33 µm min^−1^) and ∼2.75-fold lower than unmodified MTs (0.22 µm min^−1^). The dashed lines represent the 25th and 75th quartiles. The graph represents a median (±SEM) of three independent experiments with images from multiple chambers quantified from each experiment. The individual experiment data can be found in Fig. S3. **E.** Non-linear regression curve for the number of depolymerisation events of a particular rate for each of the MT variants. The distribution curve shows that most glycylated MTs depolymerised between 0.01 - 0.1 µm min^−1^ while unmodified MTs had maximum MTs with a rate between 0.2-0.3 µm min^−1^ and brain MTs had the maximum MTs between 0.3-0.6 µm min^−1^.

The other class of MAPs that regulate MT stability is the severing enzymes. Among these, katanin has been implicated in regulating cilia across species ^38,44–46^ and its activity is tightly regulated by the levels of glutamylation and glycylation of MTs ^11^ . A similar AAA-ATPase, spastin has also been implicated in regulating primary cilia dynamics in neural stem cells thus regulating neuron development, suggesting that it has a key role in cilia maintenance as well^30,47^. Considering spastin depends on glutamylation for its activity^9,10^, we wanted to understand whether glycylation has an antagonistic effect to glutamylation on spastin similar to that on katanin^11^.

Our investigations on the dependence of spastin on specific modifications (Fig. 3A) showed that similar to earlier reports^9,48^, brain MTs were severed by spastin efficiently. On the other hand, both the unmodified and glycylated MTs showed fewer severing events (Fig. 3B,C, Movie S4). The kymographs established that the severing and MT catastrophe was slower with unmodified MTs and more so with glycylated MTs (Fig. 3C). Quantifying the extent of severing revealed that brain MTs were severed much faster with 15% severing by 75 s, reaching to ∼62% by 5 min (Fig. 3D, Fig. S4, Movie S4, Table 1, Table S4). Contrary to this, the unmodified MTs as well as the glycylated MTs had very low severing with only 31 % severing observed for the unmodified MTs even after 5 min, and ∼22 % for the glycylated MTs (Fig. 3D, Fig. S4, Movie S4, Table 1, Table S4). Moreover, not only the extent of severing, but also the rate of MT severing was drastically reduced for glycylated MTs (Fig. 3E), where glycylated MTs were severed at a rate of 0.61 events µm^−1^ min^−1^, the unmodified and brain MTs were severed at a rate of 1 event µm^−1^ min^−1^ and 1.23 events µm^−1^ min^−1^ respectively. These suggest that while spastin depends on a graded level of glutamylation ^10^, higher levels of glycylation in the absence of glutamylation dramatically reduces MT severing by spastin.

**Figure 3:**
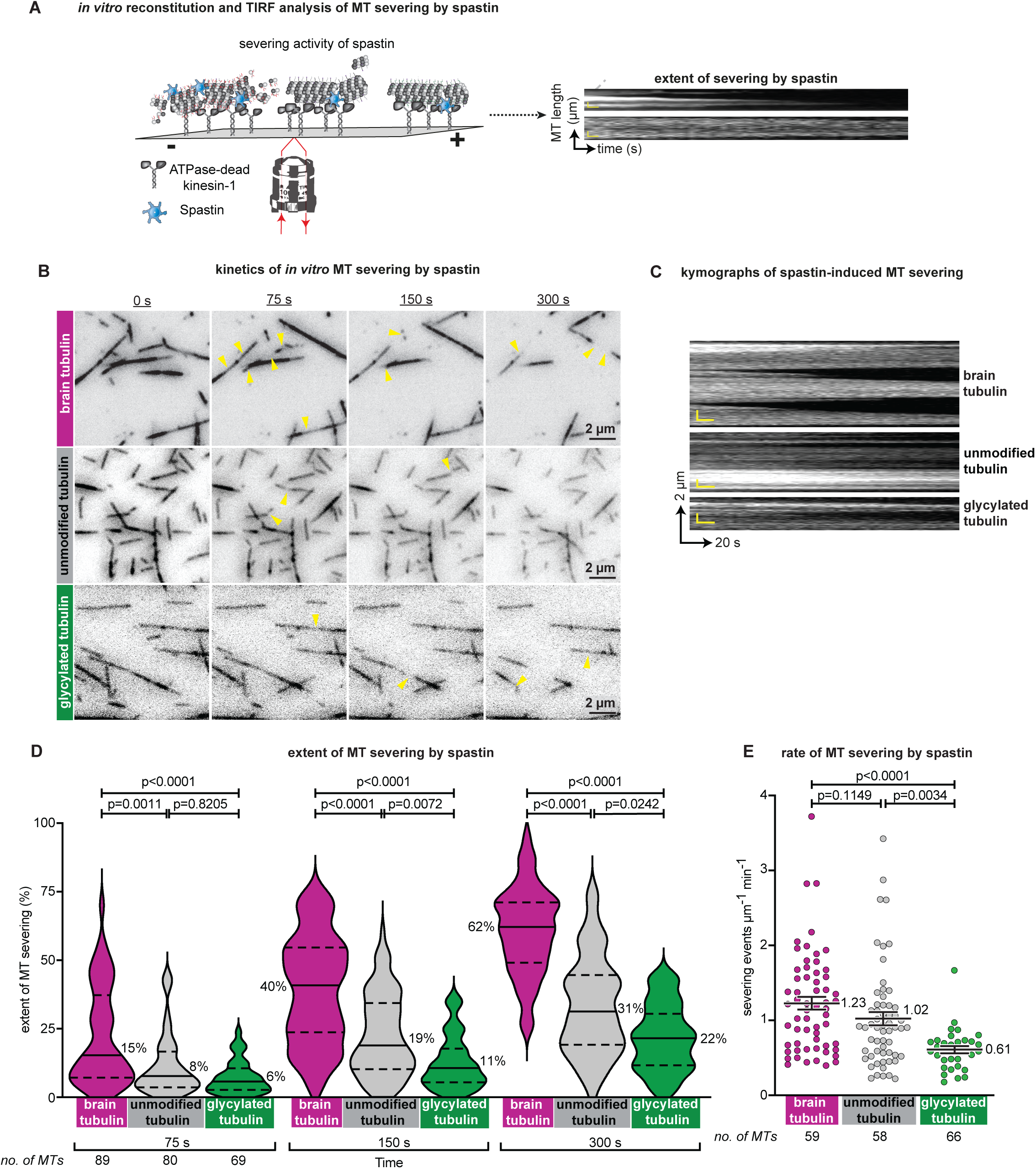
Glycylation negatively regulates spastin-induced MT severing. **A.** Schematic depiction of the TIRF microscope-based *in vitro* spastin-induced MT severing of each of the MTs with distinct PTM pattern. The imaging chambers coated with ATPase-mutant Kif5B are incubated with either of the MTs with specific PTMs or unmodified MTs, followed by introducing 8 nM spastin and ATP in the imaging motility buffer. Immediately, the MTs are visualized in TIRF to record the severing activity of spastin. Subsequently, kymographs are established and the rate of MT severing and extent of severing quantified. **B.** Snapshots of the kinetics of spastin-induced severing of the MTs with different PTM patterns. The images at 0 s, 75 s 150 s and 300 s establish that brain MTs experience rapid severing with multiple severing events observed even at 75 s. Once severed, the MTs undergo catastrophe, leading to complete depolymerisation by 300 s. The unmodified MTs and glycylated MTs showed much less severing events and extent of severing even after 300 s. Yellow arrowheads indicate the positions on the MTs undergoing severing. **C.** Representative kymographs of MTs being severed by spastin, which depict that after the severing, the MTs undergo rapid catastrophe. The kymographs reveal that among the three MTs, brain MTs show multiple severing-induced catastrophe events, while the activity and extent of severing is highly reduced for both unmodified MTs as well as the TTLL3-glycylated MTs. The x-axis depicts the time (s) and y-axis the MT length (µm). **D.** Violin plots of the quantification of the extent of spastin-induced severing shows that the extent of severing of glycylated MTs is ∼22 % even after 300 s, compared to 62 % for brain MTs. Unmodified MTs were severed to ∼31% by 300 s. Even at the earlier time point of 75 s, the glycylated MTs had ∼2.5-fold lower than brain MTs and the unmodified MTs were reduced by ∼1.3-fold. The graph represents a mean (±SEM) with the numbers and solid line denoting the median value of three independent experiments with multiple chambers imaged and quantified for each experiment. The dashed lines represent the 25th and 75th quartiles. Individual experiments are shown in Fig. S4. **E.** Scatter plot showing the rate of MT severing by spastin per µm per min, for each of the PTM variant MTs and unmodified MTs. Glycylated MTs had the slowest rate of severing with a mean of 0.61 events µm^−1^ min^−1^ which was approximately ∼1.7-fold and ∼2-fold slower than unmodified MTs (1.02 events µm^−1^ min^−1^) and brain MTs (1.23 events µm^−1^ min^−1^) respectively.

Overall, these studies show that glycylation acts to reduce the efficacy of MCAK and spastin, and as proposed earlier ^11,49^, this could potentially be one of the molecular reasons for increased stability and cilia length when glycylation is increased in cells ^17^ and why cilia shorten in the absence of glycylation ^18^.

### Motors and MAPs depend on balanced levels of glutamylation and glycylation

Our *in vitro* studies show how microtubules with only glycylation can varyingly impact certain ciliary motors and MAPs. However, both the PTMs share modification sites and manipulating one modification can influence the other ^13,18,26^. Moreover, across species cilia have been shown to always have a combination of both these PTMs (reviewed in ^1,2,49^) with glutamylation suggested to precede glycylation in multiciliated cells^35^. Hence to understand how motors and MAPs can be influenced by a combination of different levels of these modifications, we tested the same set of motors and MAPs with MTs having different proportions of glutamylation and glycylation.

To study this, apart from using brain MTs and glycylated MTs, we prepared MTs with gradients of glycylation by mixing brain tubulin and glycylated tubulin at molar ratios corresponding to final glycylation proportions of 50% and 70%. We confirmed the increasing glycylation and reducing glutamylation in these MTs by immunoblots for both the modifications (Fig. 4A) and used these four MT variants for our assays. Our analyses with Osm3 revealed that as we increased the proportion of glycylated tubulin, we observed an increase in MT gliding velocities (Fig. 4B, Fig. S5A, Movie S5, Table 1, Table S5), which increased from 257 nm s^−1^ for brain MTs to 396 nm s^−1^ for 70% glycylated MTs and 579 nm s^−1^ for glycylated MTs. This strongly suggests that the ciliary kinesin depends on glycylation for maximum activity.

**Figure 4:**
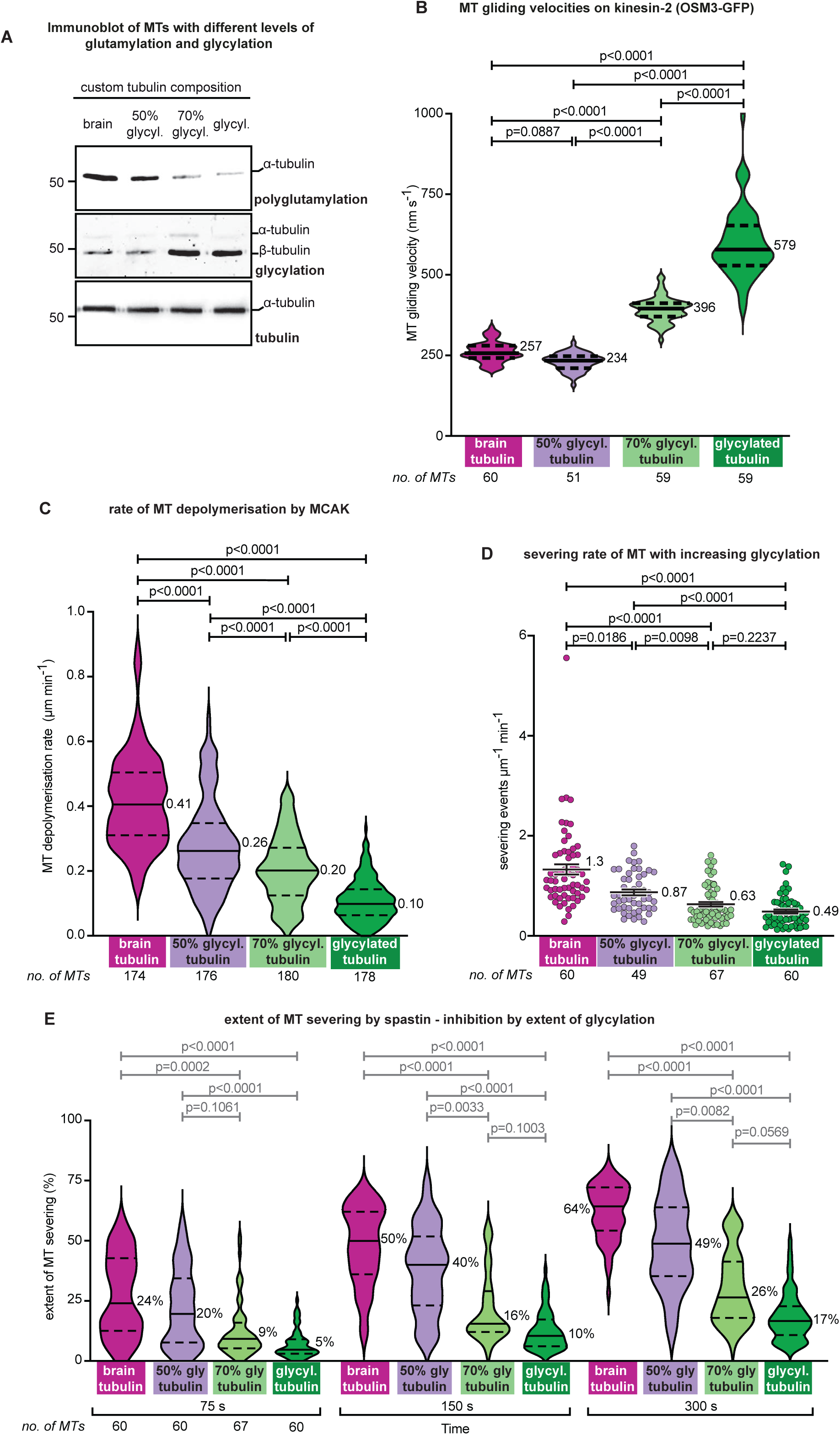
Motor and MAP activity is fine-regulated by levels of MT glutamylation and glycylation. **A.** Our custom-generated mixture of glutamylated and glycylated MTs were subjected to SDS-PAGE and Western blot analysis to confirm the levels of both the PTMs in the mixed MTs. The immunoblots for polyglutamylation and glycylation show the levels of glycylation increasing and glutamylation reducing proportionately as we titrate the molar ratios of tubulin. Blot for α-tubulin levels shows the equal tubulin load in all the lanes. **B.** Violin plot of the distribution of gliding velocities of MTs with gradient levels of glycylation as obtained from the *in-vitro* MT gliding on Osm3ΔH2. The graph shows that the velocity increased in a graded manner with increasing glycylation from a median velocity of 257 nm s^−1^ for brain MTs to 396 nm s^−1^ for 70% glycylated and 579 nm s^−1^ for glycylated MTs. The graph represents a mean (±SEM) of three independent experiments with the solid line being the median and the dashed lines representing the 25th and 75th quartiles. Individual experiments are shown in Fig. S5A. **C.** Quantification of the rate of depolymerisation of the different gradient MTs by MCAK is represented as a violin plot with the solid line and number representing the median rate and the dashed lines, the 25th and 75th quartiles. The graph highlights the gradient reduction of the depolymerisation rate with increase in glycylation in the polymerised MTs. Brain MTs, which are 90% glutamylated have a rate of 0.41 µm min^−1^ which decreases by ∼1.5-fold, ∼2-fold and ∼4-fold 50%-, 70%- and glycylated MTs. The graph is a median of three independent experiments. Individual replicates are shown in Fig. S5B. **D.** Scatter plot of the quantification of the rate of severing by spastin for the 4 different MT mixtures of glutamylated and glycylated tubulin. The plot shows that the mean rate of severing drops from 1.3 events µm^−1^ min^−1^ for brain MTs to 0.87, 0.63 and 0.49 events µm^−1^ min^−1^ for 50 % glycylated, 70 % glycylated and only glycylated MTs respectively. Graph is representing a mean (±SEM) of three independent experiments where images were acquired from multiple chambers for each experiment. **E.** Violin plot for the quantification of the severing activity of spastin after 75. s, 150 s and 300 s of imaging of each of the 4 different MT mixtures of glutamylated and glycylated tubulin. The graph highlights the gradual reduction in the extent of MT severing as the molar concentration of glycylated tubulin was increasing. The numbers and solid lines indicate the median value and the dashed lines represent the 25th and 75th quartiles. From the plots, it is evident that the extent of severing of brain MTs increases from 24% at 75 s to 64% by 300 s, while the 50 % glycylated, 70 % glycylated and only glycylated MTs show a ∼1.3-fold, ∼2.5-fold and ∼4-fold reduction in severing at every time point of imaging and quantification. The plot is a mean (±SEM) of three independent experiments with images acquired from multiple chambers for each experiment. Individual experiment data is represented in Fig. S6

We next tested the same set of MTs with MCAK and we observed that as we increased the concentration of glycylated MTs, there was a concomitant reduction in the rate of depolymerisation with 50% glycylated MTs showing a 1.5-fold reduction, 70% glycylated with 2-fold reduction and 4-fold reduction of the rate when the MTs are only glycylated (Fig. 4C, Fig. S5B, Movie S6, Table 1, Table S5). Similarly, with spastin as well, we observed that as glycylation levels increased, the severing activity reduced with 50 % glycylated MTs showing a ∼1.3-fold reduction and glycylated MTs a ∼4-fold reduction in severing by spastin compared to brain MTs (Fig. 4E, Fig. S6, Movie S7, Table 1, Table S5). The severing rates also reduced with increasing glycylation, from 1.3 events µm^−1^ min^−1^ to 0.87 events µm^−1^ min^−1^ (50% glycylated MTs), 0.63 event µm^−1^ min^−1^ (70% glycylated MTs) and 0.5 events µm^−1^ min^−1^ for only glycylated MTs (Fig. 4D, Fig. S6, Movie S7, Table 1, Table S5).

Overall, these results emphasise our earlier observations that increasing glycylation, with a concomitant reduction in glutamylation, negatively regulates most proteins except the ciliary kinesin-2, Osm3. Interestingly even at equimolar concentrations of both brain MTs and glycylated MTs, there is an impact on the activities of motors and MAPs. This could be due to some degree of glycylation inherently present in brain MTs, which also influences the overall kinetics of these MAPs.

## Discussion

Regulation of microtubule function depends on microtubule interactions with various microtubule-associated proteins that include molecular motors, severing enzymes and regulatory proteins. The tubulin code has emerged as one of the pivotal fine-modulators of these intricate molecular interactions, with tubulin posttranslational modifications playing key roles in microtubule-protein interactions. Among these, tubulin acetylation is the most studied and is implicated in diverse microtubule functions. But there is some uncertainty over how it influences the binding and transport of kinesin motors^50–53^. The other majorly studied modification, tyrosination/detyrosination influences the activity of motors and MAPs differently: kinesin-1 is tyrosination-dependent, while kinesin-2 prefers detyrosinated tubulin ^8,51^. While dynein alone is unaffected by the tyrosination status, dynein-dynactin-BICD2 requires α-tubulin tyrosination for its initial loading onto the MTs ^54^. Even the microtubule depolymerising kinesin-13, MCAK prefers tyrosinated MTs, with detyrosination suppressing its activity ^8,40,55–57^. Among the microtubule severing enzymes, spastin is unaffected by the tyrosination status of MTs ^10^, while katanin requires tyrosinated MTs for enhanced severing as its regulatory domain interacts with tyrosinated MTs ^11^.

While mammalian cilia and flagella are both acetylated and detyrosinated, the axonemes are also enriched in two competing modifications occurring on the CTTs, glutamylation and glycylation which are suggested to play key roles in fine-regulation of cilia and flagella function (reviewed in ^1^). While there is evidence suggesting that modulating glutamylation impacts IFT motors ^8,58^ and the interaction and activities of specific MAPs ^8–10^, how glycylation regulates either of them is still unclear. Here, we demonstrate that enhanced glycylation of β-tubulin CTT, with some level of glycylation on α-tubulin CTT can regulate motor proteins distinctly. Not only this, but the modification also negatively regulates spastin-induced severing. We demonstrate that this regulation depends not only on the level of glycylation, but also on the complimentary levels of glutamylation on the microtubules.

Kinesin-1, the founding member of the kinesin superfamily and KIF5B is a highly processive motor involved in transport of organelles, vesicles, mRNA and other cargos in the cell cytoplasm ^32^. Despite ubiquitously expressed, with high abundance in the neurons ^59^ it is not known to localise to cilia, thus unlikely to be exposed to glycylation. The impact of glycylation on kinesin-1 activity is inverse to that observed for kinesin-2, which is intriguing. This could be potentially due to two major possibilities: β-tubulin C-terminal tail (CTT) and the extent of its glutamylation is essential for optimal velocity of Kinesin-1 ^8^. Since TTLL3 preferentially glycylated β-tubulin CTT ^12,26^, reducing the levels of glutamylation as both modifications share common modification sites, this could lead to lesser velocity of the motor. Second, positively charged patches on kinesin-1 aids the motor’s processivity through electrostatic interactions with the negatively charged CTTs of tubulin ^31,60,61^. This suggests that the kinesin-1 prefers polyglutamylated microtubules over glycylated microtubules, as glycylation neutralises the negative charges of the Carboxy-terminal tail glutamates. Hence, the binding and the gliding rates are lower with the glycylated tubulin.

Among the different kinesins involved in cilia function (reviewed in ^62^) the kinesin-2 superfamily plays a central role in anterograde intraflagellar transport (IFT), facilitating the movement of cargo from the ciliary base to the tip. Unlike kinesin-1 which depends only on modifications of β-tubulin C-terminal tail (β-CTT), kinesin-2 is influenced by contributions from both α- and β-CTTs ^8^. While its activity is increased by glutamylation, possibly through electrostatic effects, our data suggests that glycylation has a more enhancing effect both on the binding and the velocity of Osm3. Specifically, the addition of neutral glycine residues to pre-existing glutamate side chains may decrease electrostatic interactions along the microtubule lattice, thereby potentially facilitating faster motor stepping as proposed for diverse kinesin family proteins ^63,64^.

Additionally, tyrosination of α-tubulin has a strong inhibitory effect on Osm3 activity ^8^. Considering both our unmodified and glycylated MTs differ only in their level of glycylation among all PTMs (Fig. S1E), we believe that like glutamylation, even glycylation can overcome this inhibitory effect of tyrosination leading to enhanced Osm3 motor activity and why unmodified MTs have lower binding and activity with Osm3. Moreover, it has been shown earlier that Osm3 activity is regulated by glutamylation and detyrosination ^65^, both highly prevalent PTMs in cilia. We now provide another layer of regulation of ciliary IFT motors where even short chain glycylation enhances kinesin-2 motility. Considering optimal IFT is essential for appropriate ciliary signalling in response to environmental cues, whether all these PTMs are acting in concert to provide optimal kinesin-2 activity is an interesting aspect that needs further investigation.

Cilia are dynamic organelles that undergo assembly and disassembly, dependent on cell cycle re-entry (reviewed in ^66^). Cilia disassembly is enhanced by increased levels of tubulin glutamylation or reduced glycylation ^12,35,58,67,68^. On the contrary, cilia length and stability is enhanced when glycylation is increased ^17^. Since the molecular interactions between the axonemal microtubules with diverse PTMs and the MAPs regulating ciliary stability are not well established, we focussed on testing this using two prominent MT destabilisers, the kinesin-13 motors ^29,37^ like MCAK, a well-characterised depolymerizer ^69,70^ and the MT severing enzyme, spastin ^30^. Both the proteins have been shown to be influenced by glutamylation levels ^8–10^. Glycylation was recently established to be antagonistic to glutamylation in regulating katanin-based severing ^11^. With our custom-modified tubulin variants, we broadened this understanding to MCAK and spastin and observed that like katanin, both MCAK and spastin activities are dramatically reduced in presence of glycylation. This reduction in the activity of the proteins was dependent on the levels of glycylation present within the MT sample as those with ∼10% glycylation showed maximum severing while MTs with 50% glycylation already showed a 1.5 to 1.8-fold reduction in activities. This would again be due to the possibility of a change in charge on the CTT as well as as possible change in the conformation of the CTT to a more compact, collapsed conformation ^11,71^ . This could be one possible mechanism by which glycylation controls cilia stability, which needs to be validated with *in vivo* analyses.

Interestingly, the unmodified tubulin behaves distinct to either of the modified tubulins with each of these proteins as observed earlier ^8,72^. These studies showed the need for glutamylation for optimal activity of these proteins, and we build on this to emphasize the regulatory role of glycylation in controlling the activity of the motors and MAPs, specifically in the context of regulating cilia and flagella suggesting that there is a fine regulation of these proteins by individual PTMs.

### Limitations of the study

Most of our studies have been performed with tubulin predominantly glycylated at β-tubulin, though there is some degree of glycylation also present on α-tubulin C-terminal tails. The effects we thus observe could be a combinatorial effect of glycylation of both tubulin, which is difficult to elucidate with the current custom-variant of glycylated tubulin. Second, the stoichiometries we have used for glutamylated and glycylated microtubule mixtures may not reflect those present in different types of cilia. However, the effects we observe here could still be observed with actual axonemal MTs with different levels of these PTMs, though the exact magnitudes may differ. Third, our unmodified and custom-modified tubulin isolated from HEK cells are also predominantly tyrosinated (Fig. S1E) and tyrosination enhances depolymerisation by MCAK ^8^. We do not observe such enhanced activity with HEK tubulin, probably because these MTs exhibit some degree of detyrosination. Finally, the concentrations of most of the motors and MAPs we have used are similar to what is observed *in vivo*. However, in cells, the distribution of these PTMs, the presence of other motors, MAPs, and other ciliary proteins, makes regulation more complex, thereby influencing the overall outcome of these interactions and activities.

### Potential Impact

Our *in vitro* reconstitution studies build on the expanding notion that the ‘tubulin code’ is one of the key regulators of MT-MAP interactions. We establish that glycylation, a cilia-specific PTM, plays an important role in manoeuvring the motors and MAPs on microtubules. While loss of glycylation was shown to impact axonemal dynein activity ^13^, this study highlights that

MT-protein interactions are not only influenced by loss of glycylation, but that even excessive glycylation can also affect these interactions. This establishes that glycylation needs to be finely controlled in cells for optimal ciliary function. This study further raises some interesting questions. First, kinesin-1 is required for diverse transport processes within the cells with high degree of processivity and accuracy ^32^, which depends on subtle regulation by both polyglutamylation and expression of specific tubulin isotypes^8^. However, the negative regulation by glycylation is an exciting and an interesting aspect of PTM distribution and control in cells that needs further study. Second, our *in vitro* results are confined to glycylation mediated by TTLL3, predominantly a β-tubulin specific glycylase. Whether what we observe is confined to β-tubulin-dependent activities or it is independent of which CTT is modified is also an aspect that needs further analysis by establishing MTs modified by TTLL8, the α-tubulin specific glycylase. Third, considering glycylation can overcome some of the inhibitory effects of tyrosination, suggests that there appears to be a crosstalk between these PTMs that in turn subtly regulates MT-MAP interactions, which is intriguing. These open avenues for future mechanistic studies on the roles of these two PTMs in regulating MT-MAPs within the cilia and advance our understanding of the so-far-barely explored biological roles of glycylation within the cilia.

Overall, our work establishes glycylation as a central regulatory element of the tubulin code that orchestrates ciliary microtubule function. This work lays the foundation and provides important insights into the regulation of MT-MAP interactions in cilia by tubulin glycylation, and into its potential impact on cilia stability and function.

## Acknowledgements

1. S. Gadadhar acknowledges the support from iBRIC-inStem, DBT/Wellcome Trust India Alliance Intermediate Fellowship (IA/I/22/1/506238) and the start-up research grant (SRG/2023/000847) from the Science and Engineering Research Board (SERB), Department of Science and Technology, India. M. Sirajuddin acknowledges funding support from iBRIC-inStem core grants from the Department of Biotechnology, India, DBT/Wellcome Trust India Alliance Senior Fellowship (IA/S/22/2/506502), EMBO Young Investigator Programme award, and ANRF-CRG grant (CRG/2023/005854) from the Department of Science and Technology (DST), India. S. Mullick and S. Dey are supported by the Research fellowship of the Department of Biotechnology, India. R. Ganie is supported by a CSIR-SRF and S. Mahanty is supported by the M.K Bhan Young Research Fellow program (HRD-16016/10/2025-HRD-DBT) from the Department of Biotechnology, India. We would also like to thank M. Singh (Indian Institute of Science, Bangalore, India) for insightful discussions and critical analyses.

## Conflict of interest

The authors declare no competing financial interests.

## Author Contributions

**Conceptualization:** S. Gadadhar, M. Sirajuddin

**Methodology:** S. Mullick, S. Chaya, PB. Koushik, M. Sirajuddin, S. Gadadhar **Investigation:** S. Mullick, S. Chaya, S. Dey, PB. Koushik, M. Sirajuddin and S. Gadadhar **Visualization:** S. Mullick, S. Chaya, S. Dey, R. Ganie, S. Mahanty and S. Gadadhar **Validation:** S. Mullick, S. Chaya, and S. Gadadhar

**Data curation:** S. Mullick, S. Chaya, and S. Gadadhar

**Formal Analysis**: S. Mullick, S. Chaya, S. Dey, R. Ganie, S. Mahanty and S. Gadadhar

**Supervision:** M. Sirajuddin and S. Gadadhar **Project administration:** S. Gadadhar **Resources:** M. Sirajuddin and S. Gadadhar **Software:** M. Sirajuddin and S. Gadadhar **Funding acquisition:** S. Gadadhar

**Writing – original draft:** S. Mullick and S. Gadadhar

**Writing – review & editing:** S. Mullick and S. Gadadhar with inputs from all authors

## Materials and methods

### Purification of diverse custom-modified tubulins

#### brain tubulin

Isolation of tubulin from goat brain was carried out as described earlier ^25^ . Briefly, three freshly isolated brains whose meninges and white matter were removed, were homogenized and lysed in BRB80 (80 mM PIPES/KOH pH 6.8, 1 mM EGTA, 1 mM MgCl_2_, 1 mM β- mercaptoethanol, 1 mM PMSF and 1X Protease inhibitor cocktail). The lysate was clarified by centrifugation at 5,000 g for 30 min at 4 °C (JLA 8.100, Beckmann Coulter) and the clarified supernatant was subjected to three cycles of polymerisation at 37 °C (with 33% glycerol and 0.2 mM GTP in BRB80) and depolymerisation at 4 °C (with BRB80), with subsequent centrifugation at 150,000 g for 30 min after every polymerisation-depolymerisation. The second cycle of polymerisation was performed under a high salt concentration (BRB80 containing 1 M PIPES/KOH pH 6.8, 33% glycerol, 2 mM GTP) to remove all the MAPs. The resultant tubulin free from all MAPs was used for the downstream assays.

*unmodified and custom-modified tubulin:*

Tubulin from HEK cells was purified using as per established protocol ^25,73^ . Briefly, Lenti-X cells or stable mCherry-TTLL3 Lenti-X cells were grown as a starter culture in 50 ml Dulbecco’s Modified Eagle’s Medium (DMEM; Gibco # 11965092) containing 10 % fetal bovine serum (FBS; Thermo Scientific # A5256701). Once the culture was confluent, it was inoculated into four 1 litre cultures and cultured in suspension for 14 days at 37 °C, 5 % CO_2_ on a shaker rotating at 110 rpm. Thereafter, the cells were centrifuged at 250 g for 15 min at 4 °C. The cells were washed once in phosphate buffered saline (PBS pH 7.2; Gibco # 14190144) with centrifugation at 250 g for 15 min at 4 °C. The pellet was resuspended in equal volume of lysis buffer (BRB80 with 0.1% Triton-X-100) pipetted for 30 min on ice till the entire membrane integrity was lost and the lysate was centrifuged at 150,000 g for 30 min at 4 °C. The supernatant was subjected to two cycles of polymerisation at 37 °C (BRB80 containing 33 % glycerol and 2 mM GTP) and depolymerisation at 4 °C in BRB80. The tubulin retrieved in the final supernatant was confirmed to be free of MAPs and used for the downstream assays.

### Microtubule labelling

200 μM goat brain tubulin was polymerized for 1 h at 37 °C with 2 mM MgCl_2_, 2 mM GTP, 33 % glycerol in BRB80. The polymerized tubulin was layered onto a high pH glycerol cushion (100 mM HEPES pH 8.6, 1mM MgCl_2_, 1 mM EGTA, 60 % glycerol) and centrifuged at 115,000 g at 37 °C for 40 min. The pellet, containing the MTs was resuspended in the labelling buffer (100 mM HEPES pH 8.6, 1 mM MgCl_2_, 2 mM GTP, 1 mM EGTA, 40 % glycerol containing 30 mM Alexa Fluor™ 647 NHS succinimidyl ester (Invitrogen #A20006) and pipetted once every 5 min for 20 min at 37 °C. This mix was then layered onto 2.4 ml of low pH cushion (BRB80 pH 6.8, 60 % Glycerol) supplemented with 2 mM GTP and centrifuged at 278,000 g at 37 °C for 20 min to remove any free Alexa Fluor™ 647 label. The pellet containing the labelled MTs was resuspended in BRB80 and centrifuged at 80,000 g for 10 min at 4 °C. The supernatant was collected and volume made up to 1.5 ml with BRB80 pH 6.8 containing 2 mM MgCl_2_, 1 mM GTP and 33 % glycerol and allowed to polymerize for 30 min at 37 °C. This mix was centrifuged at 80,000 g for 30 min at 37 °C. The pellet was washed with warm BRB80 and resuspended in ice-cold BRB80 for 10 min at 4 °C to allow depolymerisation. Further centrifugation at 80,000 rpm for 12 min at 4 °C retrieved the labelled tubulin in supernatant that was aliquoted in smaller volumes, snap frozen and stored at −80 °C.

### Microtubule polymerisation for in vitro assays

50 μM brain tubulin was polymerised with 1.8 µM Alexa-Fluor 647 labelled tubulin, while 10 µM of either the unmodified or glycylated tubulin were polymerized with 0.4 µM Alexa-Fluor 647 labelled tubulin in BRB80 pH 6.8 containing 2 mM GTP and 20 μM taxol (Sigma Aldrich #PHL89806) at 37 °C for 16 h to ensure less than 3 % labelled tubulin in each of the distinct variants of the MTs used in the assays.

### Purification of microtubule-associated proteins

All the plasmids for motor purification (K560-GFP, Osm3ΔH2-GFP, Pf-MCAK, mSpastin_390) were a kind gift from Prof. Sirajuddin lab, iBRIC-inStem. The plasmid for ATPase-mutant kinesin (pET28-mKif5B_mut1_N1665) and spastin (pGEX-6P-mSpastin_C389) were a kind gift from Dr. Carsten Janke, Institut Curie, Paris. The motor proteins and the severing enzyme, spastin were purified from bacterial expression system.

### purification of 10x-His tagged proteins

The ATPase-mutant kinesin (mKif5B_N1665-mut1), human kinesin-1 (K560-GFP), *C. elegans* kinesin-2 (Osm3ΔH2-GFP), and *P. falciparum* MCAK were purified as 6x-Histidine tag proteins using the HisTrap HP^TM^ (Cytiva, #17524701) affinity columns. Briefly, *E.coli* BL21 cells transformed with the respective plasmids were grown in 500 ml Luria Bertini (LB) broth containing 50 µg/ml kanamycin at 37 °C, 180 rpm till the optical density reached 0.6-0.7 nm. The cultures were subsequently induced with 0.5 mM IPTG to allow protein expression at 16 °C overnight. The cells were pelleted by centrifugation at 3500 g, 10 min, 4 °C and resuspended in lysis buffer containing protease inhibitor cocktail and 1 mM PMSF (*see Table S6 for buffer details*) and sonicated using a 5 sec ON / 8 sec OFF cycle for 3 min at 38 % amplitude to ensure proper lysis. The supernatant with the protein of interest was retrieved by centrifuging the lysed cells at 15,000g for 15 min at 4 °C and filtered through a 0.45 μm syringe filter (BioFil #FPV403030) before purification. To purify the protein, the sample was incubated for 30 min on HisTrap HP agarose column at 4 °C. The protein was allowed to bind for 30 min, non-specific proteins washed with wash buffer and the bound protein eluted using a high imidazole elution buffer (see Table S6 for more details). The eluted proteins were concentrated using protein concentrators with 10 kDa molecular weight cut off (Sigma Aldrich, #UFC901024) and desalted to remove the imidazole by centrifugation at 2800 g for 20 min at 4 °C. The concentrated, purified proteins were snap-frozen in liquid nitrogen and stored at −80 °C. To ensure that the purified motors were active, the entire purification pipeline was completed within 3 h.

For K560-GFP and Osm3ΔH2-GFP, an additional microtubule binding and release was performed to retrieve only the active proteins.

### microtubule bind-and-release to enrich active motor proteins

To enrich the active motors from the purified fractions of K560-GFP and Osm3ΔH2-GFP, the motors were subjected to microtubule bind-and-release step. First, we polymerised 100 μM of brain tubulin in BRB80 containing 2 mM GTP and 40 µM taxol for 30 min at 37 °C. The polymerised MTs were layered on 400 µl of 60 % glycerol cushion and centrifuged at 80,000 g for 10 min at 37 °C. The pelleted polymerised MTs were resuspended in 100 μl of BRB80 and mixed with 100 μl of the purified motors, incubated for 15 min at 37 °C. The mix was then supplemented with 100 mM NaCl and 10 mM ATP, layered on a 60 % glycerol cushion and centrifuged at 80,000 g for 10 min at 37 °C. The supernatant obtained post centrifugation contains the active motors, which were collected, snap frozen and stored at −80 °C.

### purification of GST-Spastin

Spastin was expressed with a GST tag at C-terminus was purified using the GSTrap column (Cytiva, #17528101). Briefly, the BL21 cells transformed with pGEX-6P-mSpastin_C389 was grown in 500 ml LB broth containing 100 µM ampicillin at 37 °C, 180 rpm in a shaker incubator till it reached an optical density of 0.6-0.7 nm. It was then induced with 0.5 mM IPTG to allow protein expression at 16 °C for 4 h. The cells were subsequently pelleted by centrifugation at 3500 g for 10 min at 4 °C and resuspended in the lysis buffer containing protease inhibitor cocktail and 1 mM PMSF (*see Table S6 for buffer details*) and sonicated using a 5 sec ON / 8 sec OFF cycle for 3 min at 38 % amplitude to ensure proper lysis. The cell lysate was centrifuged at 15,000g, for 15 min at 4°C, the supernatant collected and filtered through a 0.45 μm filter. The supernatant containing GST-spastin was allowed to bind to the column for 30 min, after which the column was washed to remove non-specific proteins and spastin eluted using a reduced glutathione-based elution buffer (*see Table S6 for details on buffer compositions*). The eluted protein was concentrated using the 10 kDa protein concentrator, desalted by centrifugation at 2800 g for 20 min at 4 °C. The purified protein was snap-frozen in liquid nitrogen and stored at −80 °C.

### Microtubule gliding assays

The microtubule gliding assay was performed as described earlier ^8^. Briefly, the purified K560-GFP or Osm3ΔH2-GFP were immobilized on motility chambers prepared on acid-washed coverslips and washed with BRB80 buffer to remove all the unbound motors. Previously polymerized, labelled microtubules flushed into the motility chambers, incubated for 5 min to allow the MTs to bind to the respective motor followed by washing with BRB80 containing 20 μM taxol and 1 mg/mL β-casein (Sigma Aldrich, #C6905). Finally, a motility mix of BRB80 containing 20 µM taxol, 1 mg/ml β-casein, 2 mM ATP and the oxygen-scavenging mix (4.5 mg/mL glucose, 0.2 mg/mL glucose oxidase (Sigma Aldrich #G2133), 80 μg/mL catalase (SRL chemicals #42168)) was added and the gliding of MTs was visualized with the Nikon Ti-2 inverted spinning disk confocal laser scanning microscope equipped with the 100x TIRF objective (oil immersion; N.A 1.49) at RT. Images were acquired with the ORCA-Flash 4.0 v3 sCMOS camera (Hamamatsu) at 100 ms exposure for 3 min with a frame interval of 2 s. The videos were further processed and analysed using Fiji v2.16.0.

### Microtubule binding assays

The motility chambers were washed once with the desalting buffer (25 mM PIPES pH 6.8, 100 mM KCl, 5 mM MgCl_2_) and the ATPase-mutant KIF5B motor was perfused into the chamber and incubated for 5 min. The different Alexa647-labelled MT variants were then flowed into the coated chambers and incubated for 5 min to allow attachment followed by washing with BRB80 buffer containing 20 μM taxol and 1mg/ml β-casein. Once the labelled MTs were visualised, the chambers were flushed with GFP-tagged motors at the desired concentrations ranging from 14-15 nmoles to 280-300 nmoles (diluted in their respective desalting buffers; *see Table S6 for buffer details*) and allowed to bind to MTs for 5-15 min at RT. Images were acquired at 500 ms for the GFP-tagged motors and 100 ms for the labelled MTs on the Nikon Ti-2 inverted spinning disk confocal laser scanning microscope equipped with the 100x TIRF objective (oil immersion; N.A 1.49) at RT using the ORCA-Flash 4.0 sCMOS camera. The images were further processed and analysed using Fiji v2.16.0.

### Microtubule depolymerisation by MCAK

The motility chambers were washed once with the desalting buffer (25 mM PIPES pH 6.8, 100 mM KCl, 5 mM MgCl_2_) for ATPase-mutant kinesin and the inactive Kif5B motor was perfused into the chamber twice and incubated for 5 min. The MTs were then flowed into the coated chambers and incubated for 5 min to allow attachment followed by washing with BRB80 buffer containing 20 μM taxol and 1mg/mL β-casein. Once the labelled MTs were visualised, the chambers were flushed with MCAK assay buffer containing BRB80 with 1mg/mL β-casein, oxygen scavenging mix, 1mM ATP and 50 nM *Pf*MCAK. Images were acquired at 100 ms exposure for 5 min with a frame interval of 5 sec on the Nikon Ti-2 inverted spinning disk confocal laser scanning microscope equipped with the 100x TIRF objective (oil immersion; N.A 1.49) at RT using the ORCA-Flash 4.0 sCMOS camera. The videos were further processed and analysed using Fiji v2.16.0.

### Microtubule severing by spastin

The motility chambers were perfused with ATPase mutant Kif5B and MTs, similar to the assay conditions used for *Pf*MCAK. The spastin severing buffer contained BRB80 with 0.1 % β-mercaptoethanol, 1mg/mL β-casein, 50 mM KCl, 1 % Pluronic F127, oxygen scavenging mix, 1 mM ATP and 8 nM GST-spastin. As soon as spastin was flushed in, the chambers were imaged at 100 ms exposure, with a frame interval of 5 sec for 5 min on the Nikon Ti-2 inverted spinning disk confocal laser scanning microscope equipped with the 100x TIRF objective (oil immersion; N.A 1.49) at RT using the ORCA-Flash 4.0 sCMOS camera. The videos were further processed and analysed using Fiji v2.16.0.

### Image Analysis and Quantification

For microtubule velocity, depolymerisation and severing rate measurements, individual microtubules were manually tracked using the segmented line tool in Fiji v2.16.0 over the microtubule undergoing such events. From the kymographs, the motor velocities, rate of depolymerisation were calculated and the graphs for velocity distribution, rate of depolymerisation were generated using GraphPad Prism version 9.0. Motor velocities were quantified from maximum intensity projections of time-lapse image sequences. The trajectory of individual microtubules (MTs) was traced using a segmented line tool, and kymographs were generated via the reslice operation. Motor velocity was calculated from the slope of the kymograph, defined by the initial and final positions along the trajectory. For depolymerisation assays, kymographs were generated using the same approach, and the MT end displaying the steepest slope, which usually corresponds to the plus-end of the MTs was used to determine the depolymerisation rate. For the MT severing, images at specific time points were analysed for the extent of MT length remaining compared to the 0 min time and this was plotted using GraphPad Prism version 9.0. P-values were calculated via one-way ANOVA using the Kruskal-Wallis non-parametric test corrected for multiple comparisons using the Dunn’s statistical hypothesis test. All the quantifications from individual experiments are listed in Tables S1-S5.

### Image and video processing

Immunoblots and SDS gels were treated with Adobe Photoshop. Intensities were adjusted uniquely in a linear manner, and no additional image treatments were performed. All images and graphs were prepared in Adobe Illustrator.

For mounting the composite videos, the still panels were prepared in Adobe Illustrator and then subsequently mounted onto the individual video compositions in Adobe After Effects. Each composition was rendered as a QuickTime movie with Adobe Media Encoder and exported as 1280X720 pixel videos with QuickTime Player v10.4.

## Notes

### Competing Interest Statement

The authors have declared no competing interest.

