## Supplementary Figures S1-S5 and Supplementary Table S6 for "Tubulin glycylation regulates microtubule-protein interactions that are key for ciliary stability and trafficking"

**Supplementary information for manuscript: Tubulin glycylation regulates MT-MAP interactions that are key for ciliary stability and trafficking**

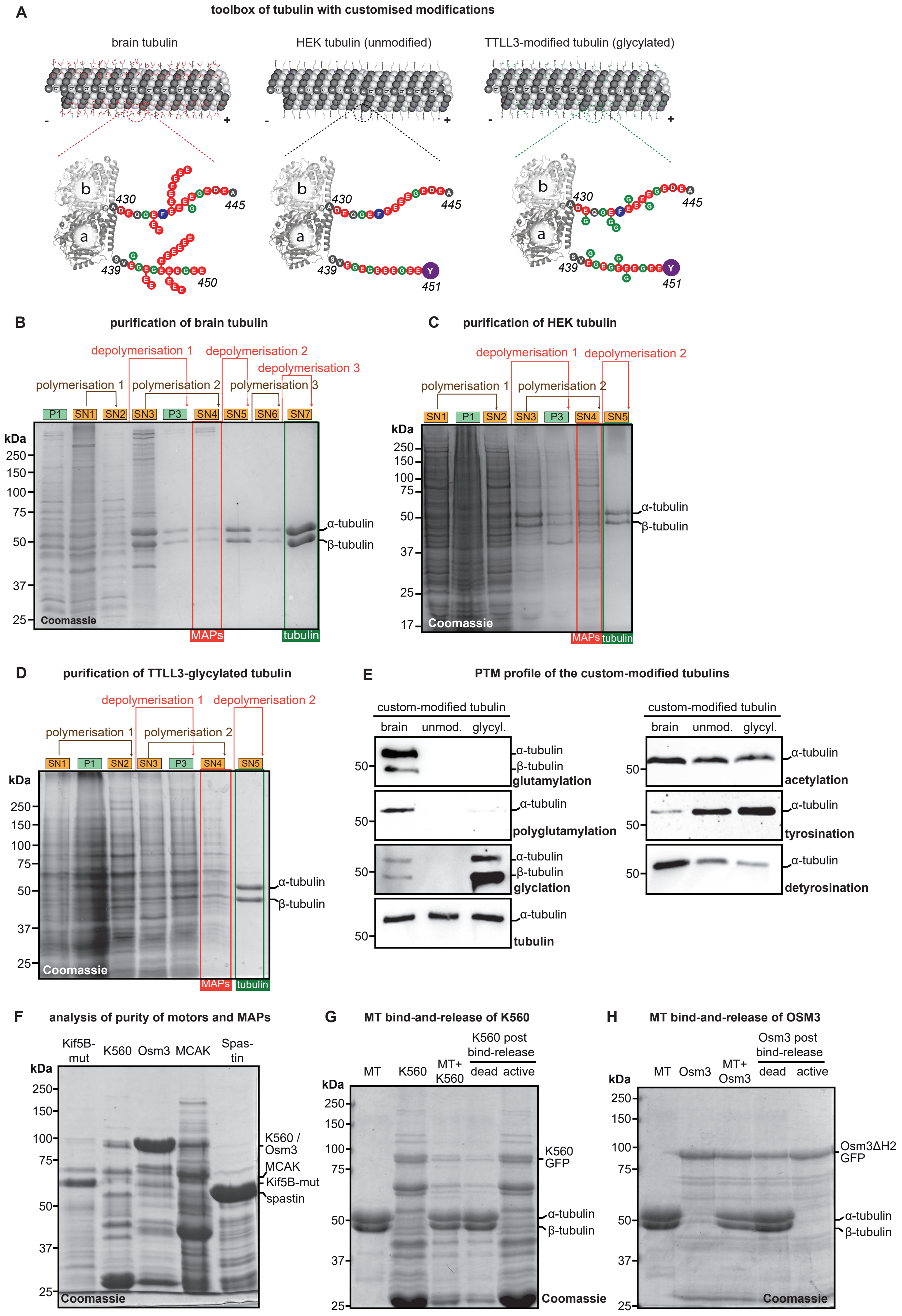
A
toolbox of tubulin with customised modifications
brain tubulin
HEK tubulin (unmodified)
TTLL3-modified tubulin (glycylated)
B
purification of brain tubulin
depolymerisation 1
depolymerisation 2
depolymerisation 3
polymerisation 1
polymerisation 2
polymerisation 3
P1 SN1 SN2 SN3 P3 SN4 SN5 SN6 SN7
kDa
250 150 100 75 50 37 25
α-tubulin
β-tubulin
Coomassie
MAPs
tubulin
C
purification of HEK tubulin
depolymerisation 1
depolymerisation 2
polymerisation 1
polymerisation 2
SN1 P1 SN2 SN3 P3 SN4 SN5
kDa
250 150 100 75 50 37 25 17
α-tubulin
β-tubulin
Coomassie
MAPs
tubulin
D
purification of TTLL3-glycylated tubulin
depolymerisation 1
depolymerisation 2
polymerisation 1
polymerisation 2
SN1 P1 SN2 SN3 P3 SN4 SN5
kDa
250 150 100 75 50 37 25
α-tubulin
β-tubulin
Coomassie
MAPs
tubulin
E
PTM profile of the custom-modified tubulins
custom-modified tubulin
brain unmod. glycyl.
α-tubulin
β-tubulin
glutamylation
polyglutamylation
glyclation
tubulin
acetylation
tyrosination
detyrosination
F
analysis of purity of motors and MAPs
Kif5B-mut K560 Osm3 MCAK Spas-tin
kDa
250 150 100 75 50 37 25
K560 / Osm3
MCAK
Kif5B-mut
spastin
Coomassie
G
MT bind-and-release of K560
MT K560 MT+K560
K560 post bind-release
dead active
kDa
250 150 100 75 50 37 25
K560 GFP
α-tubulin
β-tubulin
Coomassie
H
MT bind-and-release of OSM3
MT Osm3 MT+Osm3
Osm3 post bind-release
dead active
kDa
250 150 100 75 50 37 25
Osm3ΔH2 GFP
α-tubulin
β-tubulin
Coomassie

**Supplementary Figure S1: Toolbox of custom-modified microtubules, motors and MAPs used in the study**

**A.** Illustration of the diverse custom-modified tubulins established and analysed in the study. The schematic shows the possible levels of the glutamylation and glycylation on brain MTs, HEK MTs and TTLL3-glycylated MTs. The zoomed-in tubulin dimer shows the range of modifications in the CTT tail of α- and β-tubulin in each of these different MTs.

Coomassie blue stained SDS-PAGE gels showing the purification profile of **(B)** brain tubulin, **(C)** HEK tubulin and **(D)** TTLL3-glycylated tubulin where the gels show sequential enrichment of tubulin with each cycle of polymerization-depolymerization, finally yielding ≥95 % pure tubulin. P: pellet fraction; SN: supernatant fraction.

**E.** 2 μM of each of the tubulin variants was electrophoresed on SDS-PAGE followed by immunoblotting with PTM specific antibodies, to establish the entire profile of the different posttranslational modifications. α-tubulin was probed to assess the protein load. The blots reveal that brain MTs were enriched with glutamylation and detyrosination with very low glycylation and tyrosination. HEK tubulin, on the other hand was rich in tyrosination and low in all other PTMs corresponding to the CTT. TTLL3-modified tubulin was also heavily tyrosinated and enriched in glycylation, mainly of β-tubulin. There was no change in the level of acetylation in either of the tubulin variants.

**F.** Coomassie blue stained SDS-PAGE gels showing the purification profile of ATPase-dead Kif5B, K560-GFP, Osm3ΔH2-GFP, MCAK and spastin purified from bacteria. The multiple bands observed in the motor fraction is owing to the possible intermediate assembly fractions of the motors.

**G.** Coomassie blue stained SDS-PAGE showing the enrichment of active kinesin-1 motors following a microtubule bind-and-release assay. The motor, upon incubation with MTs, is sedimented with the MTs, which is subsequently incubated with ATP to release the active motors, which can be observed in the final lane of the PAGE. The absence of any MTs in this lane confirms that the motors collected here are the active motors, that have detached from the MTs upon ATP-driven motor activity.

**H.** Coomassie blue stained SDS-PAGE showing the enrichment of active kinesin-2 motors following a microtubule bind-and-release assay. Following the incubation with MTs, kinesin-2 sediments with the MTs, which upon treatment with ATP, is released. This fraction, observed in the final lane of the PAGE is the active kinesin-2 motor released upon ATP-driven motor activity. The absence of any MTs in this lane confirms that the motors collected here are the active motors.

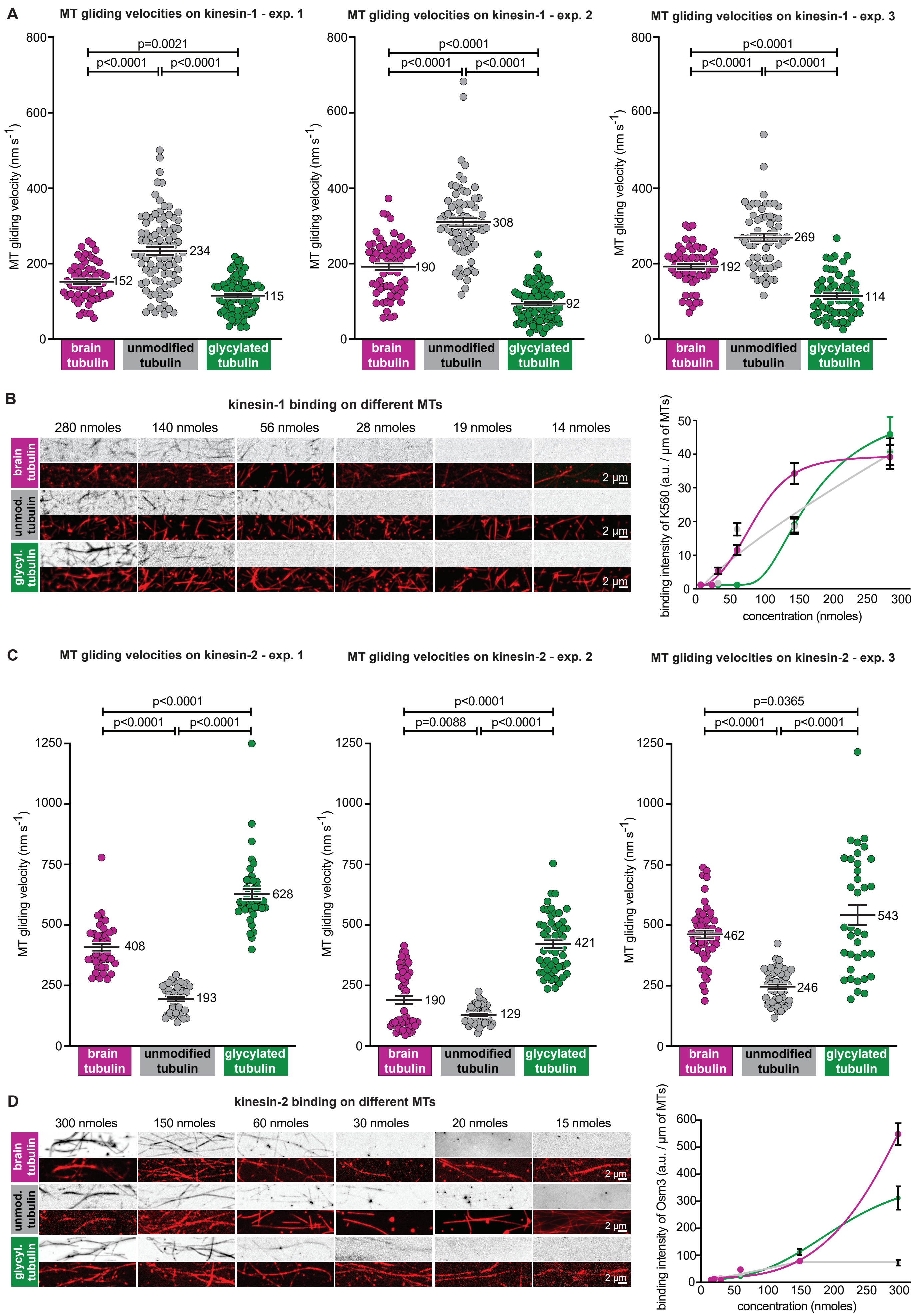
A
MT gliding velocities on kinesin-1 - exp. 1
MT gliding velocities on kinesin-1 - exp. 2
MT gliding velocities on kinesin-1 - exp. 3
MT gliding velocity (nm s-1)
p=0.0021
p<0.0001
p<0.0001
152
234
115
190
308
92
192
269
114
brain tubulin
unmodified tubulin
glycylated tubulin
B
kinesin-1 binding on different MTs
280 nmoles
140 nmoles
56 nmoles
28 nmoles
19 nmoles
14 nmoles
brain tubulin
unmod. tubulin
glycyl. tubulin
2 µm
binding intensity of K560 (a.u. / µm of MTs)
concentration (nmoles)
C
MT gliding velocities on kinesin-2 - exp. 1
MT gliding velocities on kinesin-2 - exp. 2
MT gliding velocities on kinesin-2 - exp. 3
p<0.0001
p=0.0088
p=0.0365
408
193
628
190
129
421
462
246
543
D
kinesin-2 binding on different MTs
300 nmoles
150 nmoles
60 nmoles
30 nmoles
20 nmoles
15 nmoles
binding intensity of Osm3 (a.u. / µm of MTs)

**Supplementary Figure S2: Extended data for Fig. 1B and 1D - statistical quantification of MT gliding on kinesin-1 and kinesin-2 from individual experiments**

**A.** MT gliding velocity measurements from individual experiments for different biological replicates of purification for kinesin-1, the mean of which is shown in Fig. 1B. The measurements are represented as scatter plots with a line indicating the mean (value indicated) and whiskers the SEM. p-values were calculated by one-way ANOVA. For the value of individual data point, see Table S1.

**B.** MT binding assays performed with kinesin-1. Notably, any concentration lesser than 56 nmol of K560 failed to bind glycylated MTs while maintaining substantial binding to both brain and unmodified MTs.

**C.** MT gliding velocity measurements from individual experiments for different biological replicates of purification for kinesin-2, the mean of which is shown in Fig. 1D. The measurements are represented as scatter plots with a line indicating the mean (value indicated) and whiskers the SEM. p-values were calculated by one-way ANOVA. For the value of individual data point, see Table S2.

**D.** MT binding assays performed with kinesin-2. Osm3 displayed minimal difference in binding to either brain MTs or glycylated MTs, with much lesser binding observed to unmodified MTs. These indicate that glycylation may enhances kinesin-2 (Osm3) association with MTs.

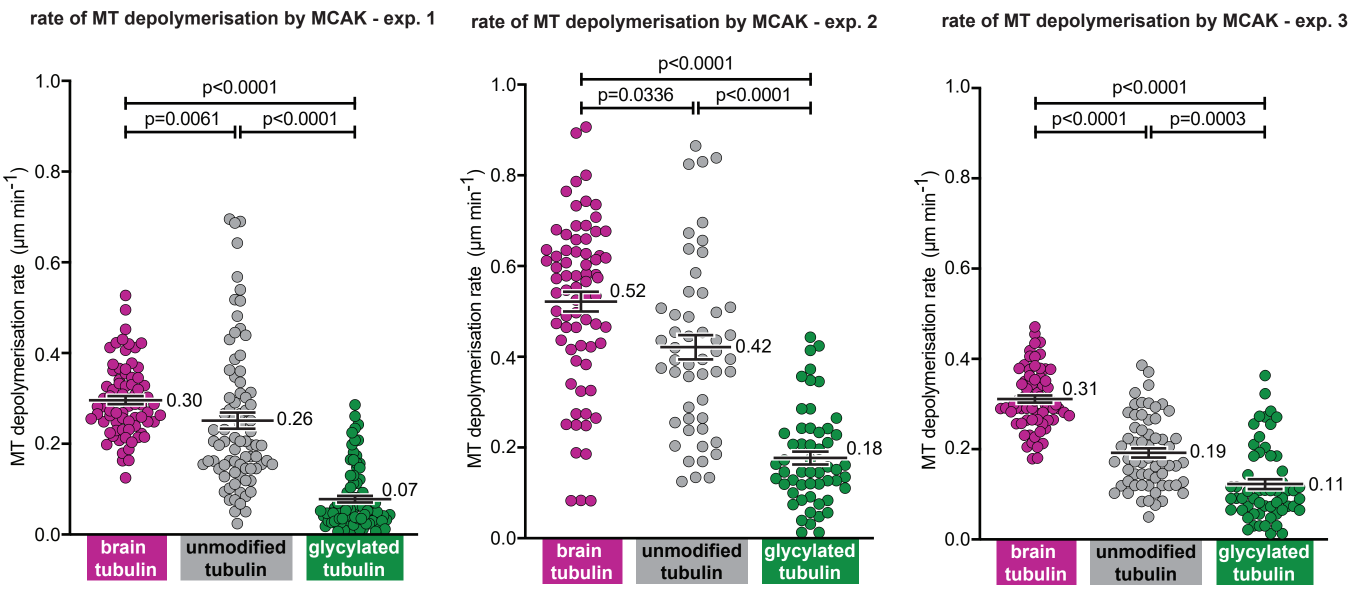
rate of MT depolymerisation by MCAK - exp. 1
rate of MT depolymerisation by MCAK - exp. 2
rate of MT depolymerisation by MCAK - exp. 3
MT depolymerisation rate (μm min-1)
1.0
0.8
0.6
0.4
0.2
0.0
p<0.0001
p=0.0061
p<0.0001
0.30
0.26
0.07
p<0.0001
p=0.0336
p<0.0001
0.52
0.42
0.18
p<0.0001
p<0.0001
p=0.0003
0.31
0.19
0.11
brain tubulin
unmodified tubulin
glycylated tubulin

**Supplementary Figure S3: Extended data for Fig. 2D - statistical analyses of rate of MT depolymerization by MCAK from individual experiments**

MT depolymerization rate from individual experiments for different biological replicates of purified MCAK. Quantification from each experiment is represented as a scatter plot of the activity of MCAK with a line indicating the mean (value indicated) and whiskers the SEM. p-values are calculated using one-way ANOVA. For the values of individual data points, see Table S3.

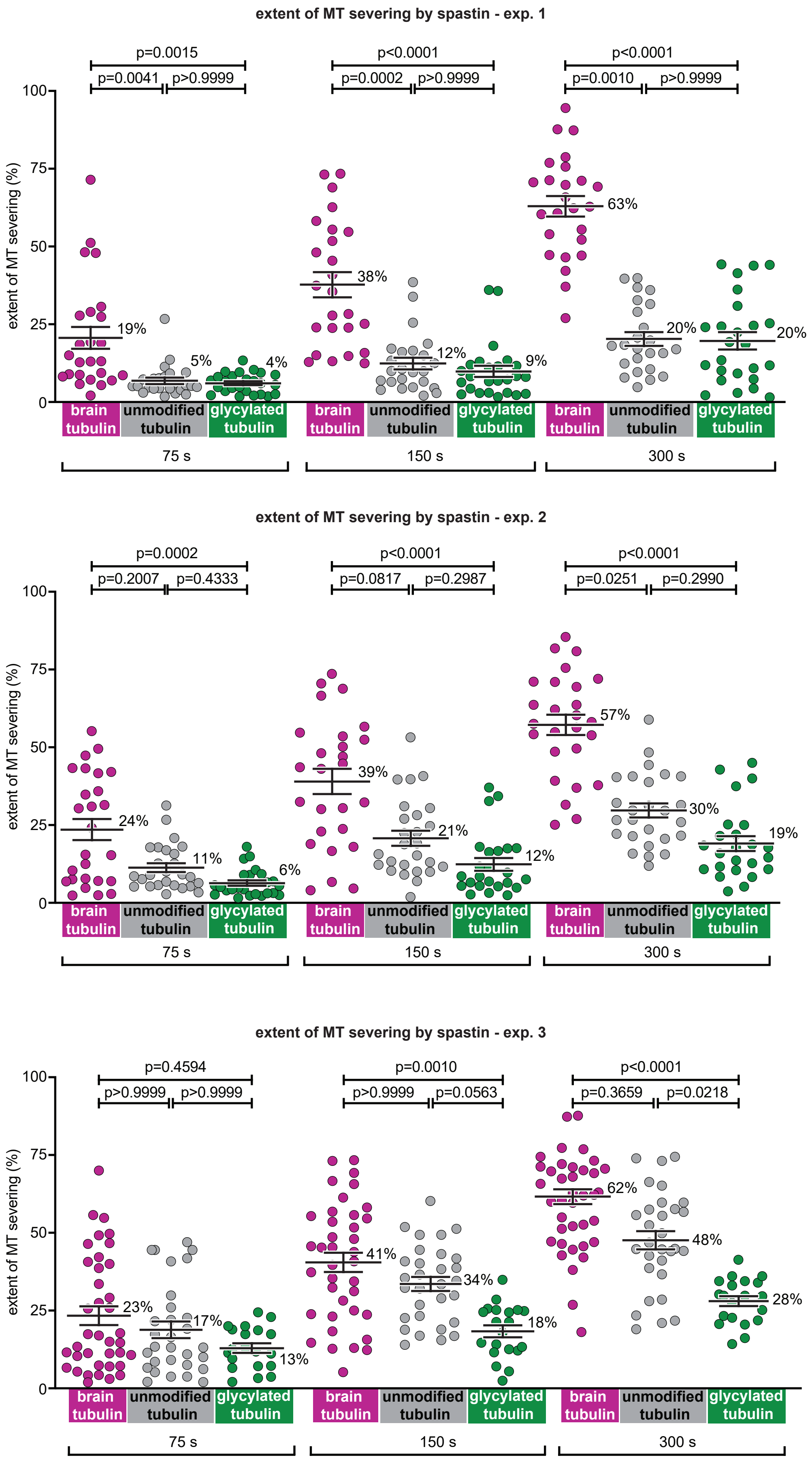

extent of MT severing by spastin - exp. 1
p=0.0015
p=0.0041
p>0.9999
p<0.0001
p=0.0002
p>0.9999
p<0.0001
p=0.0010
p>0.9999
extent of MT severing (%)
100
75
50
25
0
19%
5%
4%
38%
12%
9%
63%
20%
20%
brain tubulin
unmodified tubulin
glycylated tubulin
75 s
150 s
300 s
extent of MT severing by spastin - exp. 2
p=0.0002
p=0.2007
p=0.4333
p<0.0001
p=0.0817
p=0.2987
p<0.0001
p=0.0251
p=0.2990
24%
11%
6%
39%
21%
12%
57%
30%
19%
extent of MT severing by spastin - exp. 3
p=0.4594
p>0.9999
p>0.9999
p=0.0010
p>0.9999
p=0.0563
p<0.0001
p=0.3659
p=0.0218
23%
17%
13%
41%
34%
18%
62%
48%
28%

**Supplementary Figure S4: Extended data for Fig. 3D - statistical analyses of extent of MT severing by spastin of individual experiments**

The extent of severing of each of the MT variant by spastin is quantified for each of the individual biological replicate of purification of spastin and is represented as a scatter plot. The line indicates the mean (value indicated) and whiskers the SEM. Each graph is indicating the extent of severing across the time points of 75 s, 150 s and 300 s normalized to the extent of severing at 0 s. p-values were calculated using one-way ANOVA. For values of individual data points, see Table S4.

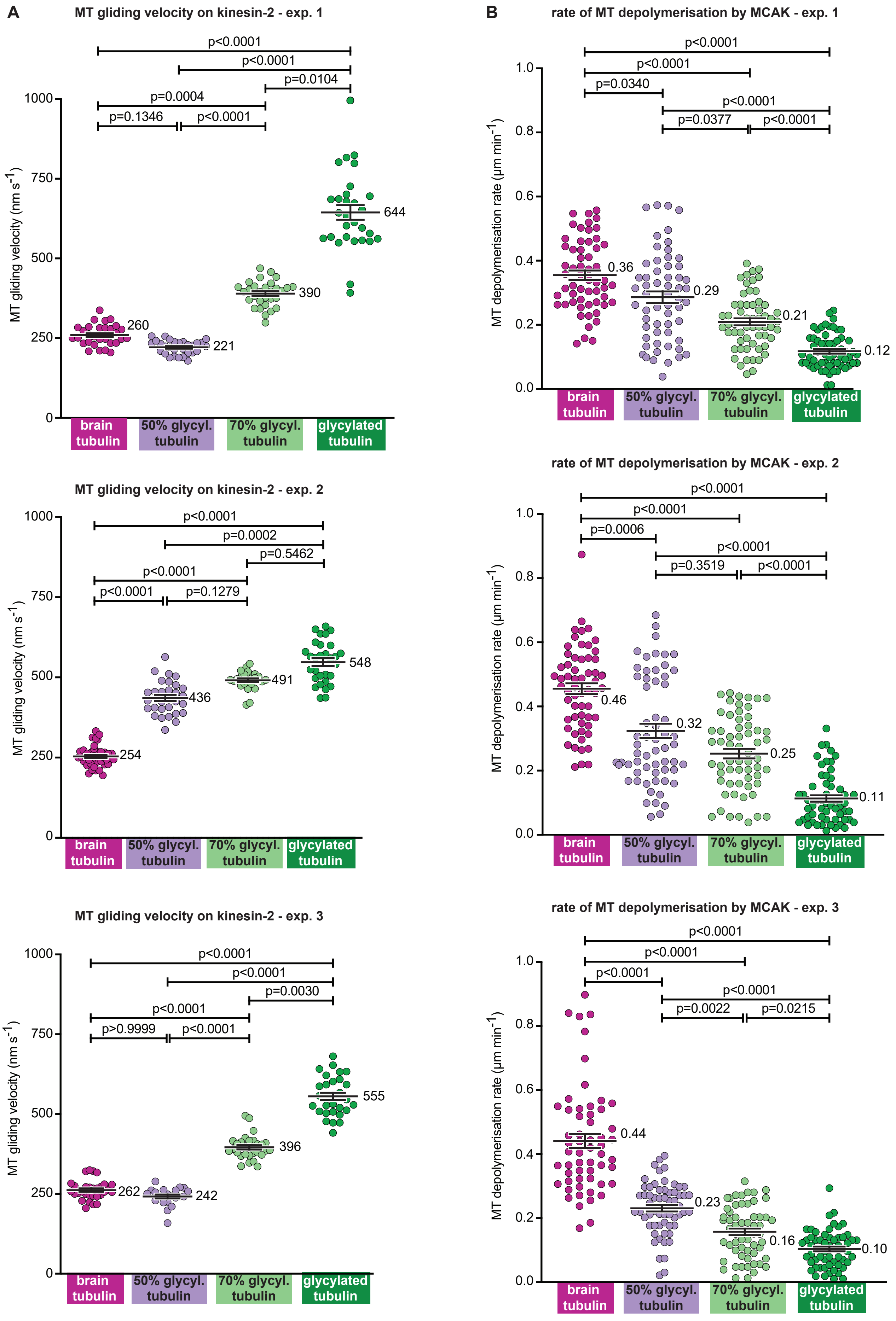
A
MT gliding velocity on kinesin-2 - exp. 1
p<0.0001
p<0.0001
p=0.0104
p=0.0004
p=0.1346
p<0.0001
MT gliding velocity (nm s-1)
1000
750
500
250
0
260
221
390
644
brain tubulin
50% glycyl. tubulin
70% glycyl. tubulin
glycylated tubulin
MT gliding velocity on kinesin-2 - exp. 2
p<0.0001
p=0.0002
p=0.5462
p<0.0001
p<0.0001
p=0.1279
MT gliding velocity (nm s-1)
1000
750
500
250
0
254
436
491
548
brain tubulin
50% glycyl. tubulin
70% glycyl. tubulin
glycylated tubulin
MT gliding velocity on kinesin-2 - exp. 3
p<0.0001
p<0.0001
p=0.0030
p<0.0001
p>0.9999
p<0.0001
MT gliding velocity (nm s-1)
1000
750
500
250
0
262
242
396
555
brain tubulin
50% glycyl. tubulin
70% glycyl. tubulin
glycylated tubulin
B
rate of MT depolymerisation by MCAK - exp. 1
p<0.0001
p<0.0001
p=0.0340
p<0.0001
p=0.0377
p<0.0001
MT depolymerisation rate (µm min-1)
1.0
0.8
0.6
0.4
0.2
0.0
0.36
0.29
0.21
0.12
brain tubulin
50% glycyl. tubulin
70% glycyl. tubulin
glycylated tubulin
rate of MT depolymerisation by MCAK - exp. 2
p<0.0001
p<0.0001
p=0.0006
p<0.0001
p=0.3519
p<0.0001
MT depolymerisation rate (µm min-1)
1.0
0.8
0.6
0.4
0.2
0.0
0.46
0.32
0.25
0.11
brain tubulin
50% glycyl. tubulin
70% glycyl. tubulin
glycylated tubulin
rate of MT depolymerisation by MCAK - exp. 3
p<0.0001
p<0.0001
p<0.0001
p<0.0001
p=0.0022
p=0.0215
MT depolymerisation rate (µm min-1)
1.0
0.8
0.6
0.4
0.2
0.0
0.44
0.23
0.16
0.10
brain tubulin
50% glycyl. tubulin
70% glycyl. tubulin
glycylated tubulin

**Supplementary Figure S5: Extended data for Fig. 4B and 4C - quantification of kinesin-2 and kinesin-13 activity on MTs with different ratios of glutamylated and glycylated tubulin**

**A.** Quantification of the activity of kinesin-2 from the individual *in-vitro* MT gliding assay for MTs with different levels of glycylation representing the experiments for different biological replicates of purification of kinesin-2. Each experiment is represented as a scatter plot with the line indicating the mean (value indicated) and whiskers the SEM. p-values were calculated using one-way ANOVA and the individual data point values can be seen in Table S5. The combined data is shown in Fig. 4B.

**B.** Quantification of the activity of MCAK from individual *in-vitro* MT depolymerization assay using different biological replicates of purification of MCAK. The data is represented as scatter plots with the line indicating the mean (value shown) and the whiskers the SEM. The combined data is shown in Fig. 4C. p-values were calculated using one-way ANOVA. Values of individual data points are found in Table S5.

**extent of MT severing by spastin of MTs with increasing levels of glycylation - exp. 1**

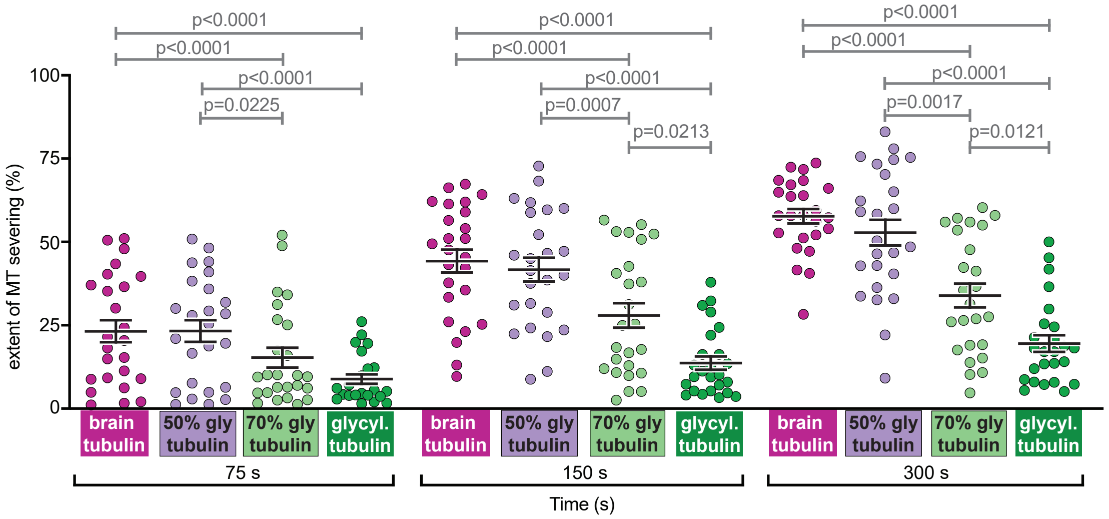

**extent of MT severing by spastin of MTs with increasing levels of glycylation - exp. 2**

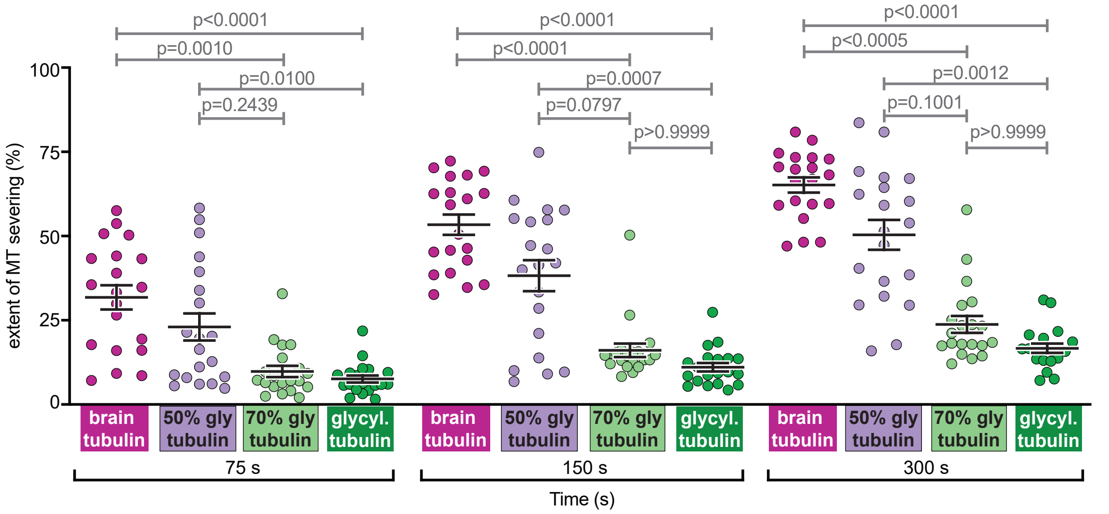

**extent of MT severing by spastin of MTs with increasing levels of glycylation - exp. 3**

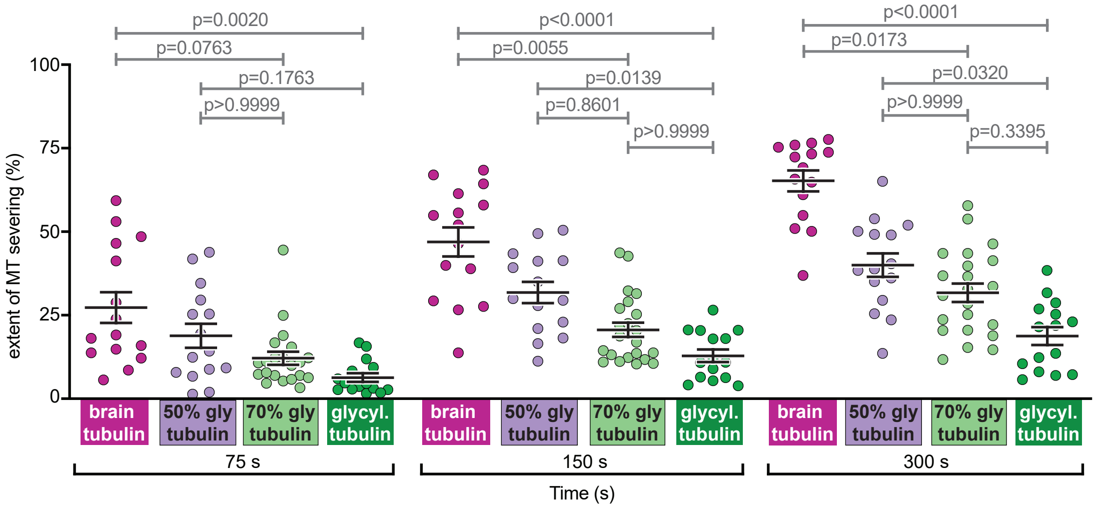

**Supplementary Figure S6: Extended data for Fig. 4D - statistical quantification of spastin activity on MTs with different ratios of glutamylated and glycylated tubulin**

Scatter plots of severing activity from individual experiments of biological replicates of purification of spastin. The line depicts the mean (value shown) and the whiskers the SEM. For each of the different MT variant, the extent of severing was determined at 75 s, 150 s and 300 s, normalised to 0 s and plotted. The combined data is shown as a violin plot in Fig. 4E. p-values were calculated using one-way ANOVA and the values for each data point are shown in Table S5.

**Supplementary Table S6: Buffer compositions for the affinity protein purifications of motors and MAPs**

| Proteins | Equilibration | Lysis | Wash A | Wash B | Wash C | Elution | Desalting |
|---|---|---|---|---|---|---|---|
| K560-GFP | 25 mM PIPES (pH 6.8)<br>400 mM KCl<br>5 mM $MgCl_2$ | 25 mM PIPES (pH 6.8)<br>400 mM KCl<br>5 mM $MgCl_2$<br>5mM β-mercaptoethanol<br>0.1% Triton<br>1 mM PMSF<br>1X PIC | 25 mM PIPES (pH 6.8)<br>400 mM KCl<br>5 mM $MgCl_2$ | 25 mM PIPES (pH 6.8)<br>400 mM KCl<br>5 mM $MgCl_2$<br>5 mM β-mercaptoethanol<br>30 mM Imidazole<br>200 μM ATP | 25 mM PIPES (pH 6.8)<br>400 mM KCl<br>5 mM $MgCl_2$<br>5 mM β-mercaptoethanol<br>50 mM Imidazole | 25 mM PIPES (pH 6.8)<br>100 mM KCl<br>5 mM $MgCl_2$<br>5 mM β-mercaptoethanol<br>250 mM Imidazole | 25 mM PIPES (pH 6.8)<br>100 mM KCl<br>5 mM $MgCl_2$ |
| Osm3ΔH2-GFP | 50 mM HEPES (pH 7.8)<br>100 mM KCl | 50 mM HEPES (pH 7.8)<br>100 mM KCl | 50 mM HEPES (pH 7.8)<br>100 mM KCl | 50 mM HEPES (pH 7.8)<br>300 mM KCl | 50 mM HEPES (pH 7.8)<br>100 mM KCl | 50 mM HEPES (pH 7.8)<br>100 mM KCl | 50 mM HEPES (pH 7.8)<br>100 mM KCl |

| | | | | | | | |
|---|---|---|---|---|---|---|---|
| | 5 mM $MgCl_2$ | 5 mM $MgCl_2$<br><br>5 mM β-mercaptoethanol<br><br>0.1% Triton<br><br>1 mM PMSF<br><br>1X PIC | 5 mM $MgCl_2$ | 5 mM $MgCl_2$<br><br>5 mM β-mercaptoethanol<br><br>30 mM Imidazole<br><br>200 μM ATP | 5 mM $MgCl_2$<br><br>5 mM β-mercaptoethanol<br><br>50 mM Imidazole | 5 mM $MgCl_2$<br><br>5 mM β-mercaptoethanol<br><br>350 mM Imidazole | 5 mM $MgCl_2$ |
| *Pf*MCAK | 25 mM PIPES (pH 6.8)<br><br>100 mM KCl<br><br>5 mM $MgCl_2$ | 25 mM PIPES (pH 6.8)<br><br>100 mM KCl<br><br>5 mM $MgCl_2$<br><br>5 mM β-mercaptoethanol<br><br>0.1% Triton<br><br>1 mM PMSF<br><br>1X PIC | 25 mM PIPES (pH 6.8)<br><br>100 mM KCl<br><br>5 mM $MgCl_2$ | 25 mM PIPES (pH 6.8)<br><br>400 mM KCl<br><br>5 mM $MgCl_2$<br><br>5 mM β-mercaptoethanol<br><br>20 mM Imidazole<br><br>500 μM ATP | 25 mM PIPES (pH 6.8)<br><br>100 mM KCl<br><br>5 mM $MgCl_2$<br><br>5 mM β-mercaptoethanol<br><br>40 mM Imidazole | 25 mM PIPES (pH 6.8)<br><br>5 mM $MgCl_2$<br><br>5 mM β-mercaptoethanol<br><br>250 mM Imidazole | 25 mM PIPES (pH 6.8)<br><br>5 mM $MgCl_2$<br><br>5 mM β-mercaptoethanol |

| | | | | | |
|---|---|---|---|---|---|
| mSpastin_C389 | 20 mM Tris-HCl (pH 7.5)<br>300 mM NaCl<br>2 mM $MgCl_2$ | 20 mM Tris-HCl (pH 7.5)<br>300 mM NaCl<br>2 mM $MgCl_2$<br>0.1% Triton<br>1 mM ATP<br>1 mM PMSF<br>1X PIC | 20 mM Tris-HCl (pH 7.5)<br>500 mM NaCl | 20 mM Tris-HCl (pH 8)<br>300 mM NaCl<br>10 mM $MgCl_2$<br>5 mM DTT<br>20 mM reduced glutathione | 20 mM HEPES (pH 7.5)<br>100 mM NaCl<br>5 mM DTT<br>3 mM $MgCl_2$ |
